# A phospho-switch provided by LRR receptor-like kinase, ALK1/QSK1/KIN7, prioritizes ABCG36/PEN3/PDR8 transport toward defense

**DOI:** 10.1101/2022.05.11.491457

**Authors:** Bibek Aryal, Jian Xia, Zehan Hu, Tashi Tsering, Jie Liu, John Huynh, Yoichiro Fukao, Nina Glöckner, Hsin-Yao Huang, Gloria Sáncho-Andrés, Konrad Pakula, Karin Gorzolka, Marta Zwiewka, Tomasz Nodzynski, Klaus Harter, Clara Sánchez-Rodríguez, Michał Jasiński, Sabine Rosahl, Markus Geisler

**Affiliations:** University of Fribourg, Department of Biology, Fribourg, Switzerland; College of Life Sciences, Ritsumeikan University, Shiga, Japan; Zentrum für Molekularbiologie der Pflanzen, Pflanzenphysiologie, Universität Tübingen, Auf der Morgenstelle 32, 72076 Tübingen, Germany; Department of Biology, ETH Zurich, Zurich, Switzerland; Department of Forest Genetics and Plant Physiology, Umea Plant Science Center, Sweden; Department of Plant Molecular Physiology, Institute of Bioorganic Chemistry, Polish Academy of Sciences, Z. Noskowskiego 12/14, 61-704 Poznań, Poland; Department of Biochemistry and Biotechnology, Poznan University of Life Sciences, Dojazd 11, 60-632 Poznań, Poland; Mendel Centre for Plant Genomics and Proteomics Masaryk University, CEITEC MU Kamenice 5, Building A26. CZ-625 00 Brno, Czech Republic; Department Biochemistry of Plant Interactions, Leibniz Institute of Plant Biochemistry, Weinberg 3, D-06120 Halle (Saale), Germany; Julius Kühn-Institut, Königin-Luise-Str. 19, D-14195 Berlin, Germany; NanoBioMedical Centre, Adam Mickiewicz University, Poznan, Poland

## Abstract

Based on its proposed substrate preferences, the ABC transporter, ABCG36/PDR8/PEN3, from the model plant Arabidopsis stands at the cross-road between growth and defence. Recently, ABCG36 was shown to export a few indolic compounds, including the auxin precursor, indole-3-butyric acid (IBA), and to be implicated in the export of the major phytoalexin of Arabidopsis, camalexin, although clear-cut proof of camalexin transport activity is still lacking.

Here we provide strong evidence that ABCG36 catalyses the direct, ATP-dependent export of camalexin over the plasma membrane, however, most likely in functional interplay with non-camalexin transporting ABCG isoforms. We identify the leucin-rich repeat receptor-like kinase, Auxin-induced LRR Kinase1 (ALK1/KIN7/QSK1), as a functional kinase to physically interact with and phosphorylate ABCG36. ABCG36 phosphorylation by ALK1 represses unilaterally IBA but not camalexin export leading to a prioritization of ABCG transport toward defense. As a consequence, phospho-dead mutants of ABCG36, like *alk1* and *abcg36* alleles, are hypersensitive toward infection with the root pathogen, *F. oxysporum*, caused by elevated fungal progression.

Our findings indicate a novel, direct regulatory circuit between a receptor kinase and an ABC transporter determining transporter substrate specificity. It appears that growth and defense balance decisions in plants are performed on the transporter level by means of a reversible phospho-switch.

## Introduction

Growth and reproduction are fundamental to the fitness of plant species. This holds especially true for annual plants, such as the model plant, *Arabidopsis thaliana*, with a limited lifespan of 8-12 weeks. However, in nature, plants as sessile organisms constantly encounter a variety of pathogens and herbivores requiring sophisticated defense mechanisms. Pathogen recognition results in two responses: pathogen-associated molecular pattern (PAMP)-triggered immunity (PTI), induced by surface-localized receptors, and effector-triggered immunity (ETI), induced by intracellular receptors (Kadota et al., 2019). The best characterized membrane-associated receptors are leucin-rich repeat (LRR) receptor-like kinases (LRR-RLK), consisting of an extracellular LRR domain, which is varying in the number of repeats and the pace of ligand perception, a transmembrane domain and a cytoplasmic kinase moiety (Boutrot and Zipfel, 2017; Couto and Zipfel, 2016; DeFalco and Zipfel, 2021; Nurnberger and Kemmerling, 2006).

A drastic physiological consequence of prolonged PIT and ETI is growth inhibition (Huot et al., 2014; Lozano-Duran and Zipfel, 2015) because resources (mainly carbohydrates) are allocated towards defense. In order to solve this dilemma, plants have evolved refined mechanisms to balance growth-defense tradeoffs (reviewed in (Huot et al., 2014)). As a misbalance between growth and defense can result in pathogen-induced decimation of a plant population, obviously, growth-defense tradeoffs have important ecological, agricultural, and economic consequences.

While most of the molecular mechanisms and players underlying growth–defense tradeoffs are unknown, it has become obvious that they are coordinately controlled by several classes of plant hormones as well as by hormone crosstalk (Huot et al., 2014; Lozano-Duran and Zipfel, 2015). Plant hormones were initially identified as growth regulators, with auxins being the first identified and probably the best-studied (Friml, 2021; Geisler, 2021). Beside the predominant active auxin, IAA (indole-3-acetic acid), indole-3-butyric acid (IBA) has recently gained more and more interest, also because it is the active ingredient in plant propagation media (Frick and Strader, 2018).

A special feature of auxins is their cell-to-cell transport, designated as polar auxin transport (PAT), which is provided by the combined action of plasma membrane-embedded auxin transporters (reviewed in Friml, 2021; Geisler, 2021; Hammes et al., 2022). An interesting finding thus was that IAA and IBA employ different transporter subsets: while IAA is predominantly transported by members of the AUX1/LAX (AUX1 (Auxin-resistant1) and like AUX1), PIN (Pin-formed) and ABCB (B-type ABC (ATP-binding cassette transporter) families; Geisler, 2021; Hammes et al., 2022), examined members of these families apparently do not transport IBA (reviewed in Damodaran and Strader, 2019; Frick and Strader, 2018). On the other hand, IBA was shown to be exported over the PM by different members of the G family of ABC transporters (ABCGs) that seem not to transport IAA (Ruzicka et al., 2010; Strader and Bartel, 2011). ABCG37/PDR9/PIS1 (Pleiotropic drug resistance9/Polar auxin transport inhibitor sensitive1), was identified to confer resistance to auxinic herbicides (Ito and Gray, 2006) and to transport auxinic compounds, including IBA and 2.4-D (Ruzicka et al., 2010). ABCG37 seems to share a redundant function with ABCG36/PDR8/PEN3 (Pleiotropic drug resistance8/Penetration resistance3; hereafter referred to as ABCG36) in regulating auxin homeostasis and plant development (Aryal et al., 2019; Ruzicka et al., 2010). Analyses of single and double mutant phenotypes suggest that both ABCG36 and ABCG37 function cooperatively in auxin-controlled plant development (Aryal et al., 2019). Unlike established IAA transporters from the PIN and ABCB families, ABCG36 and ABCG37 show a striking, overlapping outward-facing lateral localization at the root epidermis (Aryal et al., 2019; Langowski et al., 2010; Mao et al., 2016; Stein et al., 2006), where they were suggested to negatively control rootward IBA transport by lateral IBA export (Aryal et al., 2019). On the other hand, concerted IBA export out of the root has led to the concept that ABCG36/ABCG37-dependent transport of IBA mediates interactions between the root and the soil microflora (Ruzicka et al., 2010).

Originally, mutant alleles of *ABCG36* were shown to display altered responses to diverse inappropriate pathogens of Arabidopsis (Johansson et al., 2014; Kobae et al., 2006; Stein et al., 2006), suggesting an involvement of ABCG36 in nonhost resistance. *Abcg36* mutants revealed decreased extracellular accumulation of flagelin22-induced callose (Clay et al., 2009) and hyper-accumulation of flagelin22- or pathogen-elicited indole glucosinolate derivates of the PEN2 pathway (Bednarek et al., 2009; Clay et al., 2009). A striking feature of ABCG36 is its focal accumulation at the site of leaf pathogen entry, where it is thought to export defence compounds over the plasma membrane (Mao et al., 2016; Matern et al., 2019; Stein et al., 2006; Underwood and Somerville, 2013; Xin et al., 2013). ABCG36 reallocation was shown to depend on PAMPs, such as flagellin or chitin (Underwood and Somerville, 2013). On the other hand, *abcg36* was also found to own an increased sensitivity to heavy metals (Kim et al., 2007), which seems to reflect a generic transport activity of many ABC transporters.

Over the last years, several trials were conducted to identify the unknown antimicrobial and signaling molecules that are thought to be transported by ABCG36. Lu et al. (2015) identified 4-O-b-D-glucosyl-indol-3-yl formamide (4OGlcI3F) as a pathogen-inducible, tryptophan-derived and PEN2-dependent compound that over-accumulates in *abcg36/pen3* leaf tissue. A precursor of 4OGlcI3F was proposed as an ABCG36 substrate in extracellular defense against non-adapted powdery mildew pathogens, such as *Blumeria graminis* (Lu et al., 2015). In a similar approach using the oomycete, *Phytophtora infestans* as pathogen, 4-methoxyindol-3-yl-methanol (4MeOI3M) and *S*- (4-methoxy-indol-3-yl-methyl) cysteine (4MeOI3Cys) were identified as PEN2-dependent, putative ABCG36 substrates and subsequently ABCG36-dependent transport was demonstrated (Matern et al., 2019). However, both compounds did not inhibit mycelial growth of *P. infestans in vitro* but enhanced callose deposition in of seedlings treated with flagellin (Matern et al., 2019). Recently, biological evidence was provided that also ABCG34/PDR6 and ABCG36, the latter redundantly with ABCG40/PDR12, are involved in the secretion of camalexin (3-thiazol-2′-yl-indole), the predominant phytoalexin of Arabidopsis, however, biochemical proof of transport was missing (He et al., 2019; Khare et al., 2017).

ABCG36 was shown to be phosphorylated by a range of biotic and abiotic stimuli (Benschop et al., 2007; Chen et al., 2010; Kadota et al., 2019; Niittyla et al., 2007; Nuhse et al., 2007; Nuhse et al., 2004). Recently, in a phosphor-proteomic screen aiming to identify phosphorylated residues of membrane-associated proteins upon activation of an intracellular receptor, ABCG36 phosphorylation at Ser40 and Ser45, was found to be increased (Kadota et al., 2019). Interestingly, these sites were reported as flagelin22-inducible phosphorylation sites (Benschop et al., 2007) and recently already shown to be required for the function of ABCG36 in immunity against the powdery mildew, *Blumeria graminis* (Underwood and Somerville, 2017). These results suggest that ABCG36-mediated export of anti-microbial, indolic substances (such as camalexin), is activated during both PTI and ETI by protein phosphorylation to restrict pathogen colonization. However, both proof of camalexin transport by ABCG36 and the impact of ABCG36 phosphorylation on this activity is lacking.

Here we show that ABCG36 is a valid camalexin exporter that physically interacts with and is regulated by the LRR-RLK, ALK1/QSK1/KIN7. Phosphorylation of ABCG36 by ALK1 represses IBA export leading to a prioritization toward defense during infection by the fungal pathogen *Fusarium oxysporum*.

## Results

### ABCG36 is an exporter of camalexin that functionally interacts with other non-camalexin transporting ABCGs

Based on published metabolomic profiling and camalexin detoxification assays, several ABCGs were shown to be implicated in camalexin transport (He et al., 2019; Khare et al., 2017; Matern et al., 2019; Stein et al., 2006) but for none of these clear-cut transport data were provided. In order to do so we functionally expressed all suspected camalexin transporters of the ABCG family under the control of the constitutive CaMV35S (35S) promoter in tobacco (*N. benthamiana*) using agrobacterium-mediated leaf infiltration (Henrichs et al., 2012). After loading of isolated leaf mesophyll protoplasts with custom-synthesized ^3^H-camalexin (CLX), ABCG36-mediated CLX export but not that of ABCG34/PDR6 and ABCG40/PDR12 was found to be significantly different from vector control (Fig. 1A). We included also ABCG37/PDR7/PIS1, shown to act redundantly with ABCG36 in IBA transport, and ABCG35, the closest homolog of ABCG36 (81.1% sequence identity), and both were also negative.

**Figure 1:**
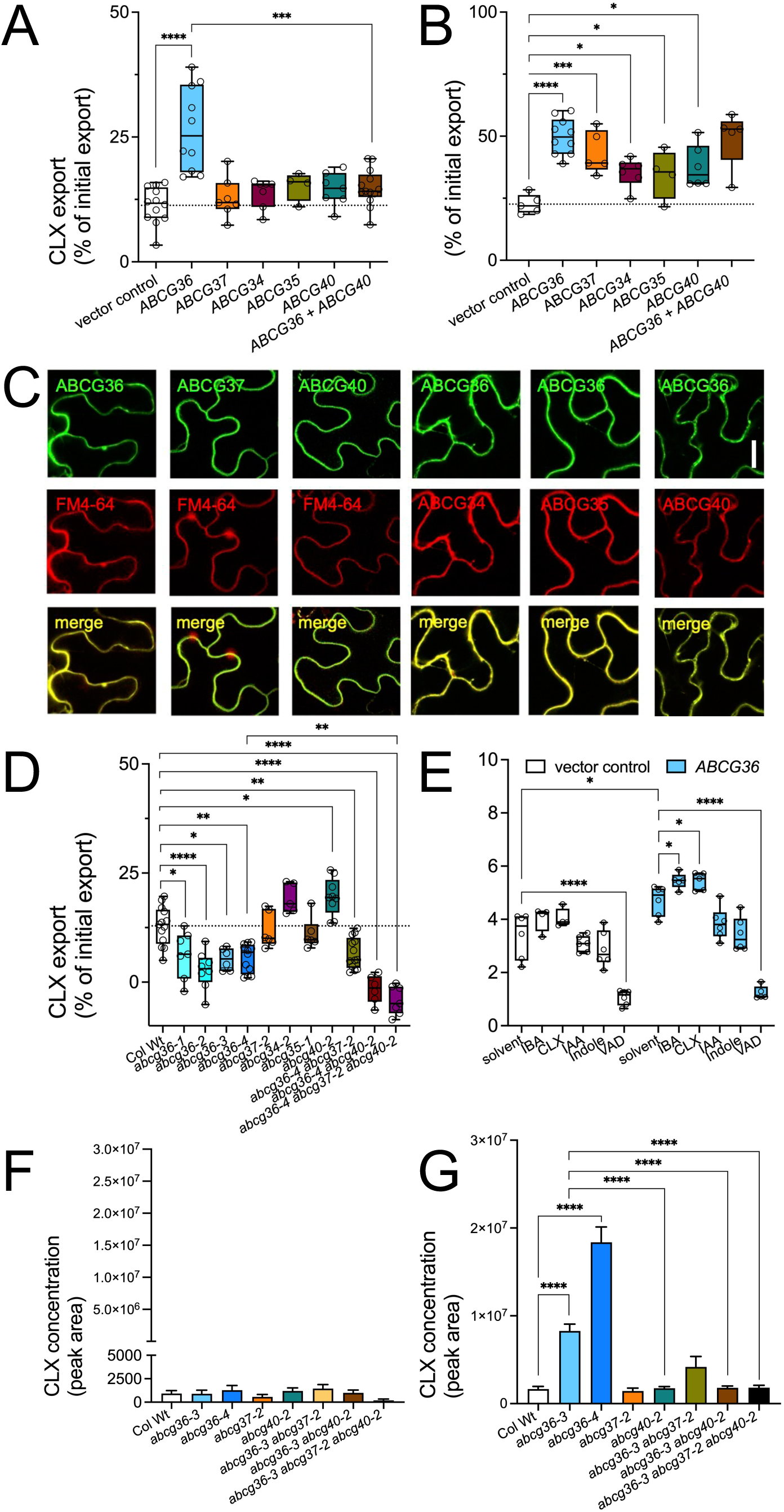
ABCG36 is an exporter of camalexin that functionally interacts with other non-camalexin transporting ABCGs. **(A-B)** CLX (**A**) and IBA (**B**) export from *N. benthamiana* protoplasts after transfection with indicated *ABCG* genes expressed under the constitutive *35SCaMV* promoter. Significant differences (Ordinary one-way ANOVA) of mean export ± SE (n ≥ 4 independent transfections) to vector control are indicated: *, *p* < 0.05; **, *p* < 0.01; ***, *p* < 0.001; ****, *p* < 0.0001). **(C)** Confocal imaging of GFP- and RFP-tagged versions of ABCGs after transfection of *N. benthamiana* leaves. FM4-64 or ABCG36-RFP were used as PM markers; bar, 50 μm. **(D)** CLX export from *Arabidopsis* protoplasts prepared from indicated *ABCG* loss-of function alleles. Significant differences (Ordinary one-way ANOVA) of mean export ± SE (n ≥ 4 independent protoplast preparations) to wild-type (Col Wt) are indicated: *, *p* < 0.05; **, *p* < 0.01; ***, *p* < 0.001; ****, *p* < 0.0001). **(E)** Vanadate (Vd)-sensitive ATPase activity of microsomal fractions prepared from tobacco leaves transfected with vector control or *ABCG36*. Significant differences (Ordinary two-way ANOVA) of mean activity ± SE (n = 3 independent microsome preparation) to vector or solvent control are indicated: *, *p* < 0.05; **, *p* < 0.01; ***, *p* < 0.001; ****, *p* < 0.0001). **(F-G)** Extracellular levels of camalexin in control (**F**) and *P. infestans*-inocculated (**G**) *Arabidopsis* leaves determined by UPLC-ESI-QTOF-MS. Significant differences (Ordinary one-way ANOVA) of mean levels ± SE (n≥ 7 with each 30 droplets) to wild type are indicated: *, *p* < 0.05; **, *p* < 0.01; ***, *p* < 0.001; ****, *p* < 0.0001).

We tested the same set also for IBA transport and discovered that all tested ABCGs were capable of exporting IBA, although ABCG34, G35 and G40 to a lesser amount than ABCG36 and G37 (Aryal et al., 2019; Ruzicka et al., 2010; Fig. 1B). Importantly, all tested ABCGs were correctly and to similar levels expressed on the PM as could be judged by colocalization with PM markers, FM4-64 and ABCG36-GFP, respectively (Fig. 1C).

In order to back-up heterologous expression data, we quantified CLX export from protoplasts prepared from Arabidopsis *ABCG* loss-of-function mutants. In agreement with tobacco transport data, all four tested *abcg36* alleles but not the single *abcg35* and *g37* mutant alleles showed significantly reduced CLX transport compared to the wild type (Wt; Fig. 1D). Interestingly, in contrast to tobacco transport Arabidopsis *abcg34* and *abcg40* lines revealed significantly increased export rates compared to the Wt.

To further support this apparent dual transport activity for IBA and CLX on ABCG36, we performed ATPase and competition assays. Vanadate-sensitive ATPase activity, which can be attributed to ABC transporters (Locher, 2008; Verrier et al., 2008), was significantly increased for microsomes prepared from tobacco transfected with ABCG36 in comparison to the vector control (Fig. 1E). More importantly, ABCG36-specific ATPase activity was significantly increased by its substrates, IBA and CLX, a hallmark for ABC transporters including ABCGs (Hao et al., 2020; Kowal et al., 2021), but not by non-transported but structurally close, IAA or indole (Fig. 1E). In agreement, a 1.000-times access of non-labelled IBA and CLX competed-out for ATP-dependent ^3^H-IBA uptake into *ABCG36* in comparison with the solvent control but not vector control microsomes (Suppl. Fig. 3C), indicating that both substrates use most likely the same translocation pathway on ABCG36.

In order to better understand the discrepancy between the absence of transport capacities for ABCG40 and its described roles in CLX transport, we quantified transport of ABCG36 upon co-expression with ABCG40 in tobacco. Surprisingly, co-expression reduced CLX but not IBA export to vector control level (Fig. 1A, B) without changing ABCG36 expression or localization (Fig. 1C), suggesting an inhibitory role of ABCG40 on ABCG36. In Arabidopsis, CLX export of *abcg36 abcg40* was further reduced in comparison to *abcg36* alone. This effect was specific for ABCG40 as CLX export was not further altered in *abcg36 abcg37* or *abcg36 abcg37 abcg40* (Fig. 1D). Finally, we quantified exported CLX in the *abcg* accessions during nonhost resistance response of *Arabidopsis* to *P. infestans* infection by established drop-inoculation of leaves (Matern et al., 2019). As described previously, water controls of Wt showed very low surface levels of CLX (Matern et al., 2019) that were not significantly different from *abcg* loss-of-function mutants (Fig. 1F). However, surface drops including *P. infestans* zoospores contained roughly 1.800-fold enhanced CLX in Wt compared to the non-infected control, while CLX concentrations collected from *abcg36-3* and *abcg36-4* leaves were 5- and 11-fold elevated in comparison to the infected Wt, respectively (Fig. 1G). This increase is similar but stronger as previously found for the *abcg36-1/pen3-1* mutant allele (Matern et al., 2019). In agreement with our transport data, *abcg40* did not show any significant difference to Wt (Fig. 1G), however, both *abcg36 abcg37* and *abcg36 abcg37 abcg40* reduced the amount of exported CLX to WT level.

In summary, these data indicate that ABCG36 is a functional CLX exporter. However, other non-CLX transporting ABCGs, such as ABCG34 and ABCG40, seem to contribute to CLX transport by either interfering with ABCG36-mediatedCLX transport, thus acting as a negative regulator of ABCG36 or ABCG36 might function itself as regulator of these under infection (for details, see Discussion).

### ABCG36 physically interacts with the LRR receptor-like kinase, ALK1/QSK1/KIN7, *in planta*

Having on one hand the apparent dual role of ABCG36 as transporter in growth (IBA) and defense (CLX) and on the other, its putative regulation by protein phosphorylation in mind, we were speculating that both events might by controlled by a regulatory kinase. In order to identify such one, we employed a co-immunoprecipitation approach followed by tandem mass spectrometry (MS-MS) analyses as described before (Henrichs et al., 2012; Zhu et al., 2016) using microsomal fractions from *ABCG36-GFP* (*PEN3:PEN3-GFP*) roots as starting material. Three independent co-immunoprecipitation/MS-MS analyses identified a short list of 62 common, putative ABCG36 interacting proteins (Suppl. Table 1). We selected the LRR-RLK, At3g02880, for further analyses based on the following: first, At3g02880 was previously pulled-down as an ABCG36/PEN3/PDR8 interacting protein (Campe et al., 2016). Second, At3g02880 (QSK1/KIN7) was shown to regulate the potassium channel, TPK1 (Chen et al., 2021; Isner et al., 2018) and the aquaporin, PIP2;4 (Wu et al., 2019), by protein phosphorylation. Interestingly, *qsk1* mutants revealed delayed lateral root development as well as callose production suggesting an involvement in both growth and defense programs. And finally, auto-phosphorylation was found to be stimulated by auxin (Chen et al., 2010) and thus At3g02880 was named <u>A</u>uxin-stimulated <u>L</u>RR <u>K</u>inase1 (ALK1).

In order to sustain a physical interaction on the PM, we co-transfected leaves of *N. benthamiana* with *ABCG36-GFP* (*PEN3:PEN3-GFP*) and *ALK1-mCherry* (*35S:ALK1-mCherry*). Confocal imaging revealed that ABCG36-GFP and ALK1-mCherry co-localize on the PM of tobacco pavement cells (Figs. 2A, 4C). Co-immunoprecipitation using microsomes prepared from co-transfected *tobacco* allowed to detect ALK1-mCherry in the elution fraction using ABCG36-GFP but not free GFP as a bait (Suppl. Fig. 1B). Reversely, we were also able to co-IP ABCG36 from ALK1-YFP Arabidopsis lines (Suppl. Table 2).

**Figure 2:**
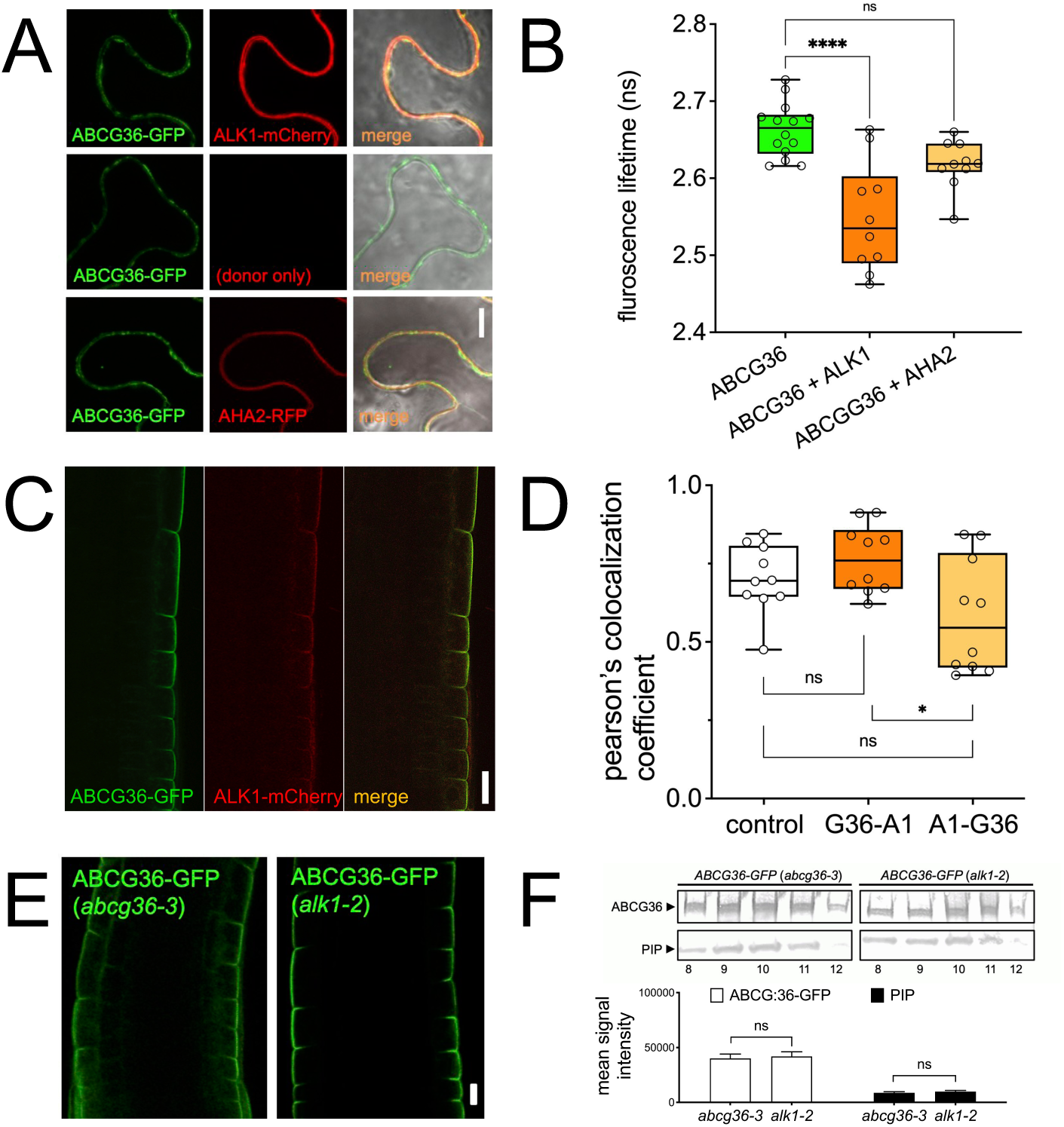
ABCG36 interacts physically and functionally with the LRR receptor-like kinase, ALK1/QSK1/KIN7. **(A)** Confocal imaging of ABCG36-GFP (PEN3:PEN3-GFP) co-transfected with ALK1-mCherry (35S:ALK1-mCherry) or AHA2-GFP (AHA2:AHA2-GFP) in *N. benthamiana*; bar, 10 μm. **(B)** FRET-FLIM analysis of ABCG36-GFP (PEN3:PEN3-GFP)/ALK1-mCherry (35S:ALK1-mCherry) interaction in *N. benthamiana* pavement cells after co-transfection. Significant differences (Ordinary one-way ANOVA) of mean fluorescence lifetime ± SE (n ≥ 10 images) to ABCG36-GFP alone are indicated: *, *p* < 0.05; **, *p* < 0.01; ***, *p* < 0.001; ****, *p* < 0.0001). **(C-D)** Confocal imaging (**C**) and colocalization coefficient analysis by Pearson’s (PC) method and Mander’s overlap for ABCG36-GFP with ALK-mCherry (G36-A1) and ALK-mCherry with ABCG36-GFP (G36-A1) in the columella and epidermis of Arabidopsis roots (**D**). Above-threshold Pearson’s correlation coefficients measured from individual confocal cross sections are indicated in a boxplot. The values represent average of at least 35 measurements. Ordinary one-way ANOVA indicates statistically significant differences (ns, *p* < 0.05); bar, 50 μm. Controls are in Suppl. Fig. 1A. **(E-F)** Confocal imaging (**E**) and Western analyses (**F**) of ABCG36-GFP (PEN3:PEN3-GFP) expressed in *abcg36-3/pen3-3* and *alk1-2*. PIP2;1 found in sucrose fractions 8-12 is used as PM marker. Significant intensity differences of quantified ABCG36 or PIP2;1 signals from peak band 10, respectively. (Ordinary one-way ANOVA) of mean signal ± SE (n = 3 independent Western analyses) between *abcg36-3/pen3-3* and *alk1-2* are indicated: ns, *p* > 0.05); bar, 50 μm.

Excitingly, ABCG36-GFP and ALK1-mCherry remarkably well co-localized in the epidermis of Arabidopsis roots as was found by crossing of stable Arabidopsis *PEN3:PEN3-GFP* and *35S:ALK1-mCherry* lines (Fig. 2C; Suppl. Fig. 1A). Despite the fact that *ALK1* was expressed under the CaMV35S promoter, ALK1 expression and colocalization with ABCG36 was limited to outer lateral domains of the epidermis. Finally, FRET-FLIM analysis of ABCG36-GFP (*PEN3:PEN3-GFP*)-ALK1-mCherry (*35S:ALK1-mCherry*) interaction in *N. benthamiana* pavement cells after *A. tumefaciens*-mediated co-transfection resulted in significantly reduced fluorescence lifetime for the ABCG36-GFP/ALK1-mCherry pair compared to the donor (ABCG36-GFP) alone, while the negative control ABCG36-GFP/AHA2-RFP did not (Fig. 2A-B). Reduced fluorescence lifetime is indicative for a very close proximity of both partners (< 10 nm) that is usually interpreted as a physical interaction. Taken together, co-localization, co-IP and FRET-FLIM analyses in tobacco and Arabidopsis, respectively, strongly support a physical interaction between ABCG36 and ALK1 on the PM.

To exclude that ABCG36-ALK1 interaction affects ABCG36 expression and/or localization as shown for some auxin transporters (Hammes et al., 2022), we crossed ABCG36-GFP (*PEN3:PEN3-GFP*) lines into the *alk1* background. Confocal imaging of roots (Fig. 2E) and Western analyses of PM fractions isolated by sucrose gradient centrifugation and verified using the PM marker, PIP2;1 (Fig. 2F), indicated no significant differences between Wt and *alk1*. These data were further hardened by immunolocalization of ABCG36/PDR8 in Wt and *alk1* using anti-PDR8 (Suppl. Fig. 1C; (Kobae et al., 2006)).

Numerous ABC transporters, including some plant hormone transporters have been shown to be up-regulated by their own substrates (Biala et al., 2017; Geisler et al., 2005; Hwang et al., 2016; Kretzschmar et al., 2011). For ABCG36, we found that low μM concentrations of IBA and CLX but not of IAA and indole significantly up-regulated ABCG36-GFP fluorescence and thus its expression in the root (Fig. 3A-B, D). Interestingly, the same behavior was found for ALK1 tested in the same lines although to a slightly lesser extend (Fig. 3A-B).

**Figure 3:**
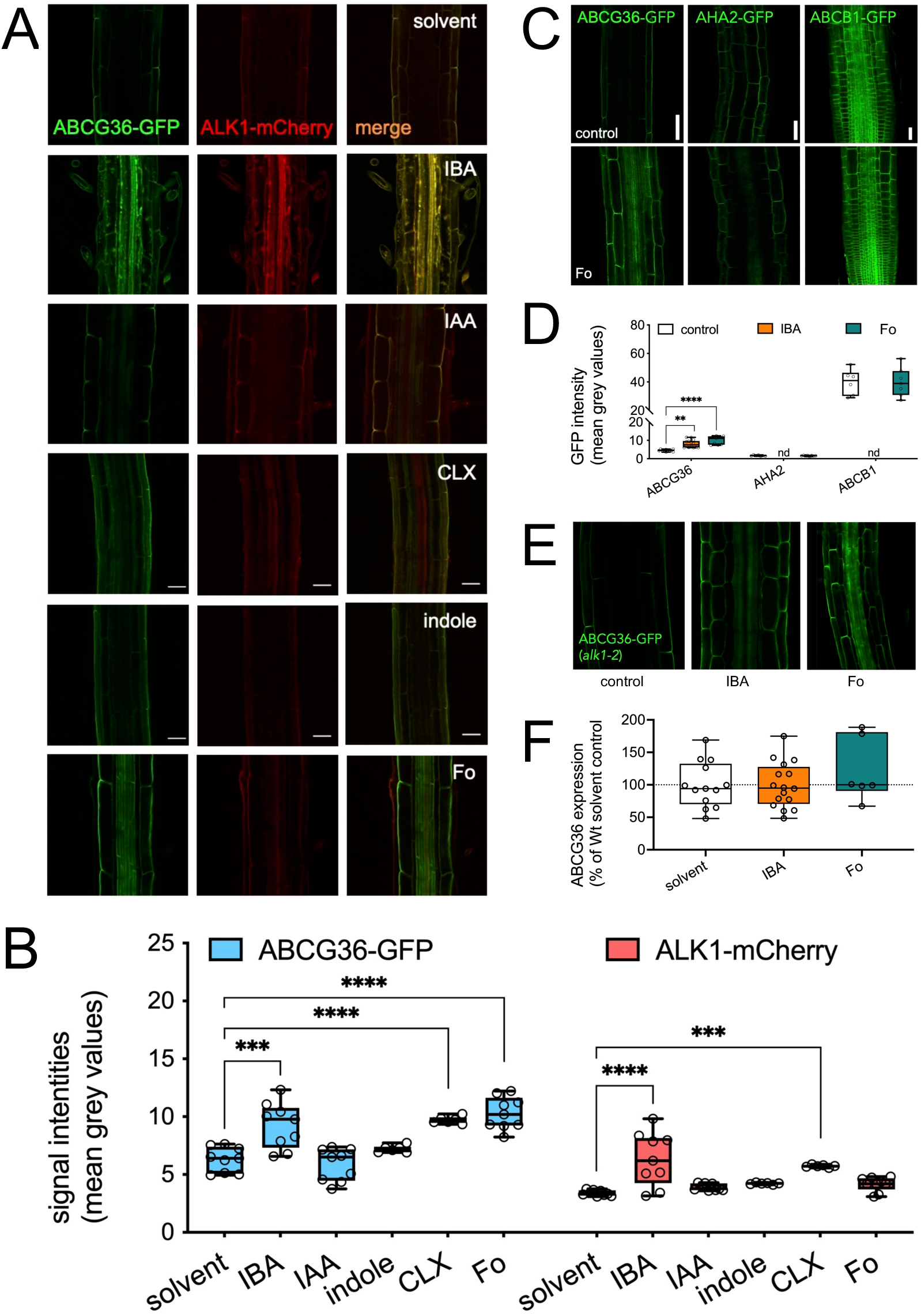
Root infection and ABCG36 substrates specifically alter ABCG36 expression and polarity. **(A-B)** ABCG36-GFP (PEN3:PEN3-GFP) and ALK1-mCherry (35S: ALK1-mCherry) was imaged **(A)** in the Arabidopsis root after indicated treatments (1 μM, 24h) in comparison with the solvent control (solvent); bar, 50 μm. Quantification of GFP and mCherry intensities **(B)**. Significant differences (Ordinary one-way ANOVA) of mean intensities ± SE (n =3 independent treatments, ≥ 50 images) to solvent control are indicated: ***, *p* < 0.001; ****, *p* < 0.0001). **(C-D)** ABCG36-GFP (PEN3:PEN3-GFP), AHA2-GFP (AHA2:AHA2-GFP) and ABCB1-GFP (ABCB1:ABCB1-GFP) was imaged 24h after treatment with *F. oxysporum* 699 (Fo) spores in comparison with buffer control (control); bars, 50 μm (**C**). Quantification of GFP intensities **(D)**. Significant differences (Ordinary one-way ANOVA) of mean GFP intensities ± SE (n =3 independent treatments, ≥ 50 images) to solvent control are indicated: **, *p* < 0.01; ***, *p* < 0.001; nd, not done. **(E-F)** ABCG36-GFP (PEN3:PEN3-GFP) was imaged in roots of *alk1-2* 24h after IBA or *F. oxysporum* 699 (Fo) treatment; bars, 50 μm (**E**). Differences (Ordinary one-way ANOVA) of mean GFP intensities in *alk1-2* ± SE (n =3 independent treatments, ≥ 50 images) are not significant from GFP intensities in *abcg36-3/pen3* solvent control (*p* > 0.05). Controls are in Suppl. Fig. 1C.

**Figure 4:**
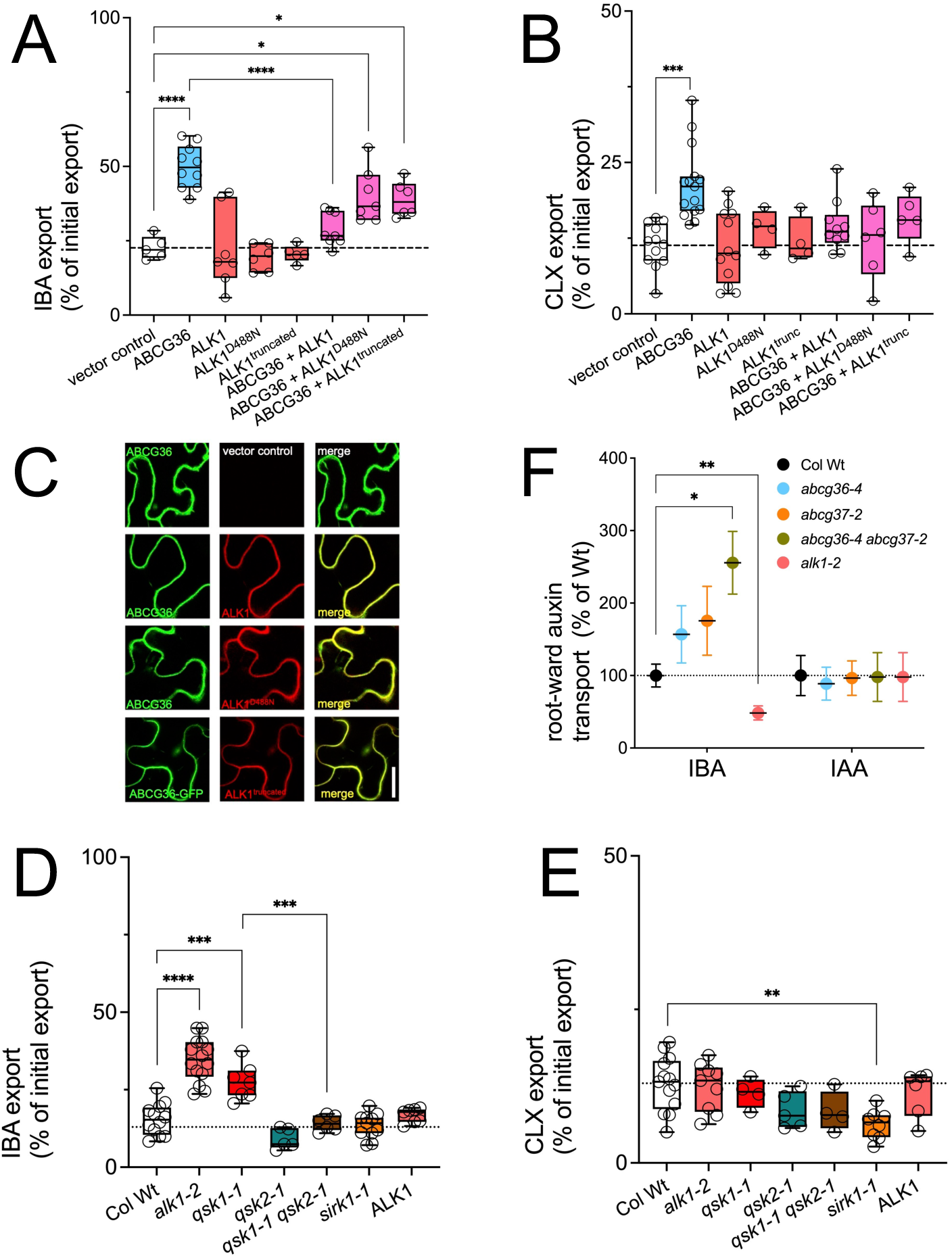
ALK1/QSK1/KIN7 is a negative regulator of ABCG36-mediated IBA efflux. **(A-B)** IBA (**A**) and CLX export (**B**) from *N. benthamiana* protoplasts after co-transfection of ABCG36 (35S:ABCG36) with Wt ALK1 (35S:ALK1-mCherry), kinase-dead ALK1 (35S:ALK1^D488N^-mCherry) or a kinase-truncated ALK1 (35S:ALK1^truncated^-mCherry). Significant differences (Ordinary one-way ANOVA) of mean export ± SE (n ≥ 4 independent transfections) to vector control or ABCG36 alone are indicated: *, *p* < 0.05; **, *p* < 0.01; ***, *p* < 0.001; ****, *p* < 0.0001). **(C)** Confocal imaging of ABCG36-GFP after co-transfection of *N. benthamiana* leaves with vector control, Wt ALK1 (35S:ALK1-mCherry), kinase-dead ALK1 (35S:ALK1^D488N^-mCherry) or kinase-truncated ALK1 (35S:ALK1^truncated^-mCherry); bar, 50 μm. **(D-E)** IBA (**D**) and CLX (**E**) export from *Arabidopsis* protoplasts prepared from indicated *LRR-RLK* loss-of function alleles. Significant differences (Ordinary one-way ANOVA) of mean export ± SE (n ≥ 4 independent protoplast preparations) to wild-type (Col Wt) are indicated: **, *p* < 0.01; ***, *p* < 0.001; ****, *p* < 0.0001). Controls are in Suppl. Fig. 3. **(F)** Root-ward IBA and IAA transport in indicated Arabidopsis Wt and loss-of function alleles. Significant differences (Ordinary one-way ANOVA) of mean transport ± SE (n = 3 independent transport assays) to wild-type (Col Wt) are indicated: **, *p* < 0.01; ***, *p* < 0.001; ****, *p* < 0.0001).

ABCG36 was shown to accumulate at the site of leaf pathogen entry (Stein et al., 2006; Xin et al., 2013) but the effect of pathogen infection on ABCG36 root abundance has not been tested yet. At first, we tested an elicitor prepared from *Fusarium oxysporum* (Fo), a well-characterized, ubiquitous root vascular fungal pathogen that causes wilt disease on several plant species, including *Arabidopsis thaliana* (Gordon, 2017). Moreover, we had previously shown that *agcg36/pen3* is hypersensitive to Fo infection (Mao et al., 2017b). Like IBA and CLX, Fo elicitor treatment resulted in strong accumulation of ABCG36 but not that of ALK1 (Fig. 3A-B). Second, we directly tested Fo infection on Arabidopsis plants grown in hydroponics revealing that this fungus, like IBA (or CLX) but even stronger, altered ABCG36 abundance (Fig. 3C-D). The effect caused by Fo was specific for ABCG36 as it was not found for other PM proteins like the H^+^-ATPase isoform, AHA2, or the auxin exporter, ABCB1 (Fig. 3C-D). Remarkably, IBA, Fo elicitor treatments and Fo infection (24h) induced ABCG36 presence in non-epidermal cell files, like the cortex and the stele. Moreover, in all cells ABCG36 loses its lateral polarity and owns widely a non-polar PM distribution. A quantification of ABCG36-GFP signals revealed a ca. 3-4 and 10-fold accumulation in the epidermis and stele upon Fo infection, respectively (Suppl. Fig. 2A). The latter was independent of transcription as revealed by quantitative real-time PCR analyses. While typical wounding markers, like transcription factors MYB51 and WRKY11, were up-regulated this was not the case for *ABCG36* transcripts, or *ABCG37, AHA2* or *NIP5;1*, employed as negative control (Suppl. Fig. 2B). In order to test the idea that ALK1 is involved in ABCG36 accumulation, we analysed ABCG36 (*PEN3:PEN3-GFP*) presence upon IBA treatment and Fo infection (24h) in the *alk1-2* background. Accumulation of ABCG36 by IBA and Fo in *alk1* was qualitatively and quantitatively not significantly different to Wt (Fig. 3E-F) making an involvement of ALK1 in these events unlikely.

In summary it appears that both ABCG36 and ALK1 are interacting proteins that are both up-regulated by valid ABCG36 substrates, IBA and CLX, on the post-transcriptional level, supporting a potential functional interaction. However, ABCG36 PM location and substrate induction is independent of ALK1.

### IBA- but not CLX- export by ABCG36 is negatively regulated by the LRR receptor-like kinase, ALK1/QSK1/KIN7

Several LRR-receptor kinases were shown to directly regulate PM transporters, including H^+^-ATPases (Fuglsang et al., 2014; Haruta et al., 2014; Li et al., 2021), Ca^2+^-ATPases (Frei dit Frey et al., 2012) and aquaporins (Wu et al., 2019). Of special interest was that ALK1/QSK1/KIN7 was shown to regulate the potassium channel, TPK1 (Chen et al., 2021; Isner et al., 2018) and the aquaporin, PIP2;4, by protein phosphorylation (Wu et al., 2019). Physical (Fig. 2) and a putative functional (Fig. 3) ABCG36-ALK1 interaction, as well as indication for ABCG36 regulation by phosphorylation (Kadota et al., 2019; Underwood and Somerville, 2017), a well-known mode of ABC transporter regulation (Conseil et al., 2001; Geisler and Hegedus, 2020; Henrichs et al., 2012), prompted us to investigate the impact of ALK1 on ABCG36 transport activity. First, we quantified ABCG36-mediated IBA efflux upon ALK1 co-expression in the tobacco protoplast system, where ALK1 expression alone had no significant effect on IBA export (Fig. 4A). However, ALK1 co-expression strikingly reduced ABCG36-catalyzed IBA export to vector control level and this inhibitory impact by ALK1 was dependent on its kinase activity: a putative kinase-dead mutant (ALK1^D488N^; (Bojar et al., 2014)) or a truncated version of ALK1 (ALK1^trunc^) lacking the entire kinase domain (cut-off after aa 286) were unable to significantly inhibit ABCG36 (Fig. 4A). Importantly, co-expression with Wt or mutated ALK1 in tobacco did not alter expression or location of ABCG36 (Fig. 4C). Interestingly, ABCG36-mediated CLX transport was slightly but not significantly inhibited by ALK1 co-expression (Fig. 4B).

Second, and in order to substantiate these transport data *in planta*, we analyzed IBA and CLX export prepared from the independent *alk1/qsk1/kin7* alleles, *alk1-2 and qsk1-1* (Wu et al., 2019). Interestingly, both alleles showed enhanced IBA but not CLX (Fig. 4D-E) or IAA (Suppl. Fig. 3A) export rates in comparison to the Wt, while ALK1 over-expression had no significant effect indicating that ALK1 activity in Wt might be in saturation (Fig. 4D-E). We included in our analyses also Sucrose-Induced Receptor Kinase1 (SIRK1), which was found to regulate aquaporins through phosphorylation under conditions of external sucrose supply by interaction with ALK1/QSK1/KIN7, as well as QSK2, a close homolog of ALK1/QSK1 (Wu et al., 2019). None of the single mutants showed a significant reduction in IBA or CLX export, however, the *qsk1 qsk2* double mutant reduced enhanced IBA export in *qsk1/alk1* to Wt level, indicating a putative involvement in ABCG36 regulation (Fig. 4D-E).

Interestingly, also *alk1* mutant roots showed an inverse behavior in root-ward/acropetal IBA transport in comparison to *abcg36 abcg37* (Fig. 4F). In line with an inhibitory action for ALK1 and a lateral, epidermal removal of IBA by ABCG36/37, absence of ALK1 thus seems to lead to enhanced export of IBA out of the root, which would dampen polar IBA streams. This regulatory impact by ALK1 was not found for IAA (Fig. 4F) underlining ALK1 specificity. In agreement, *alk1* was also lacking the hyper-sensitivity of *abcg36* (and *abcg37*) mutants upon IBA treatments visualized using the DR5:GFP responsive element (Suppl. Fig. 3B). In summary, this subset of data indicates that ALK1 negatively regulates ABCG36-mediated IBA - but not CLX - transport in an action that requires its kinase activity.

### ALK1/QSK1/KIN7 is an active kinase involved in ABCG36 phosphorylation

All validated phosphorylation sites on ABCG36 identified so far (Kadota et al., 2019; Underwood and Somerville, 2017) are concentrated in two clusters preceding the nucleotide binding domains (NBDs; Fig. 5A, Suppl. Fig. 4A). In order to identify ALK1-specific phosphorylation sites on ABCG36 that are of relevance for transport regulation, we employed a label-free, quantitative phospho-proteomic approach. Using Columbia Wt material, we were able to detect under our conditions safely four ABCG36 phosphorylation sites, S40, S45, S823 and S825, two of each cluster. Importantly, phosphorylation of S823 and S825 was strongly reduced in the *alk1* background (Fig. 5B; S37 peptides could not be detected), suggesting that these are *bona fide* ALK1 phosphorylation sites. Further, we quantified phosphorylation of these residues upon IBA, CLX and Fo5176 elicitor treatments and upon Fo infection. Strikingly, and similar as in *alk1*, phospho-sites S823 and 825 were strongly dephosphorylated by IBA in comparison to solvent control, while CLX treatment upregulated S45 but had only a mild effect on the others (Fig. 5B). Interestingly, both Fo elicitor mix and infection strongly up-regulating ABCG36 expression, were oppositely affecting ABCG36 phosphorylation: while Fo infection reduced strongly ABCG36 phosphorylation similar to IBA treatments, the effector enhanced S40 and S45 but reduced S823 phosphorylation (Fig. 5B).

**Figure 5:**
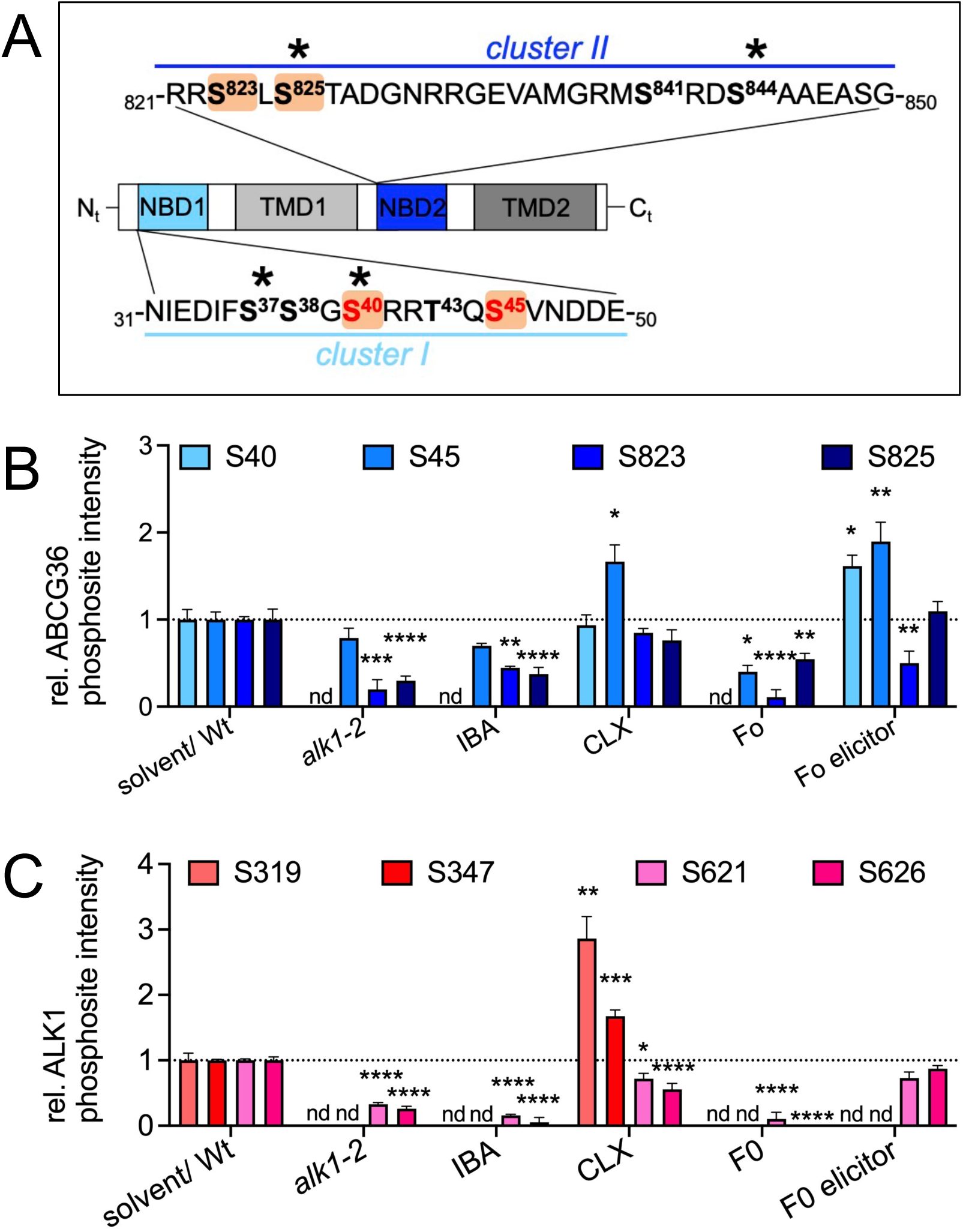
Identification of ALK1/QSK1/KIN7-specific phosphorylation sites on ABCBG36. **(A)** Proteomically supported ABCG36 phosphorylation sites identified in this study align with previously classified clusters (Underwood and Somerville, 2017) in cytosolic stretches prior to NBD1 and NBD2. Phosphorylation sites that were previously identified by phosphor-proteomics are in bold, that were previously shown to be functionally relevant during infection are colored red, that are shown here to be ALK1-specific are underlaid orange, while phospho-sites that are investigated here in more detail are marked by asterisks. NBD, nucleotide binding domain; TMD, transmembrane domain. **(B-C)** Mean abundance of indicated ABCG36 (**B**) and ALK1 (**C**) phospho-sites identified by label-free phospho-proteomics of Arabidopsis Wt (Col Wt) or *alk1-2* grown in liquid cultures. Wt treated with IBA, CLX or Fo elicitor (Fo5176 ; 1 μM, 24h) or infected with Fo (Fo699 ; 10^7^ conidia/ml; 24h). Significant differences (Ordinary one-way ANOVA) of mean phospho-site abundance ± SE (n =3 independent microsomal preparations) to solvent control or to wild type, respectively, are indicated: *, *p* < 0.05; **, *p* < 0.01; ***, *p* < 0.001; ****, *p* < 0.0001). Raw data and quality controls are in Suppl. Table 3.

Using the same datasets, we were able to detect also phosphorylation on ALK1 that likewise revealed also four phosphorylated serine residues on two clusters (Fig. 5C; Suppl. Fig. 4A). Interestingly, these phospho-sites showed overall a similar behavior to those on ABCB36 - dephosphorylation by IBA and Fo infection and enhanced phosphorylation by CLX (Fig. 5C) - supporting again a functional interconnection.

The data so far identified a clear role for ALK1 in ABCB36 phosphorylation but did not provide evidence that ABCG36 phosphorylation is directly catalyzed by ALK1. In fact, previous analyses caused mixed reports on the activity of ALK1/QSK1/KIN7 as protein kinase: while Isner et al. (2017) proposed a direct impact of ALK1/KIN7 in phosphorylation of TPK1, Wu et al. (2019) reported that PIP2;1 phosphorylation was provided by SIRK1. In this model, ALK1/QSK1 was supposed to own a regulatory role and to act as a classical co-receptor being trans-phosphorylation by SIRK1 (Wu et al., 2019). In support of the latter concept, ALK1/QSK1/KIN7 had been annotated as an atypical or pseudo-kinase based on the lack of a series of conserved residues in their kinase domain, including a phosphorylated aspartate (Castells and Casacuberta, 2007; Paul and Srinivasan, 2020). Interestingly, also SIRK1 despite being shown to be an active kinase had been annotated as a pseudo-kinase (Paul and Srinivasan, 2020). In order to clarify this issue, we performed *in vitro* kinase assays with GST-purified kinase domains of ALK1/QSK1/KIN7 and SIRK1 (Wu et al., 2019) using a peptide library of either mouse or human proteins. MS analyses identified 183/291 (mouse/human) and 93/225 (mouse/human) ALK1/QSK1/KIN7 and SIRK1-specific phospho-peptides, respectively, that showed overlapping consensus motifs (Suppl. Fig. 4C). Next, we repeated *in vitro* kinase assays with synthetic peptides containing S38 and S825 of putative cluster I and II. LC-MS/MS analyses clearly identified S38 as being specifically phosphorylated by both kinases in an ATP-dependent manner (Suppl. Figure 5A), while S825 was also partially phosphorylated in the -ATP control (Suppl. Figure 5B). These data indicate ALK1 and SIRK1 are independently active as protein kinases and that both are capable of phosphorylating cluster I phosphorylation sites of ABCG36.

### Phosphorylation of ABCG36 by ALK1/QSK1/KIN7 inhibits IBA but not CLX transport *in planta*

To demonstrate the potential impact of the phosphorylation status of individual, identified serines, we selected two from each cluster, S37, S40 and S825 and S844, respectively, and analyzed IBA and CLX transport of phospho-mimicry (S-to-D) and phospho-dead (S-to-A) *ABCG36* mutations (Fig. 5A). In agreement with ALK1 co-expression, phospho-mimicry of all four serines reduced ABCG36-mediated IBA export from transfected tobacco protoplasts to vector control level (Fig. 6A, Suppl. Fig. 6A). Excitingly, this what not the case for CLX export where none of the D mutations significantly affected ABCG36 transport (Fig. 6B, Suppl. Fig. 6A). Importantly, all four phosphor-mimicry mutations did not significantly alter ABCG36 expression or PM location (Suppl. Fig. 6B).

**Figure 6:**
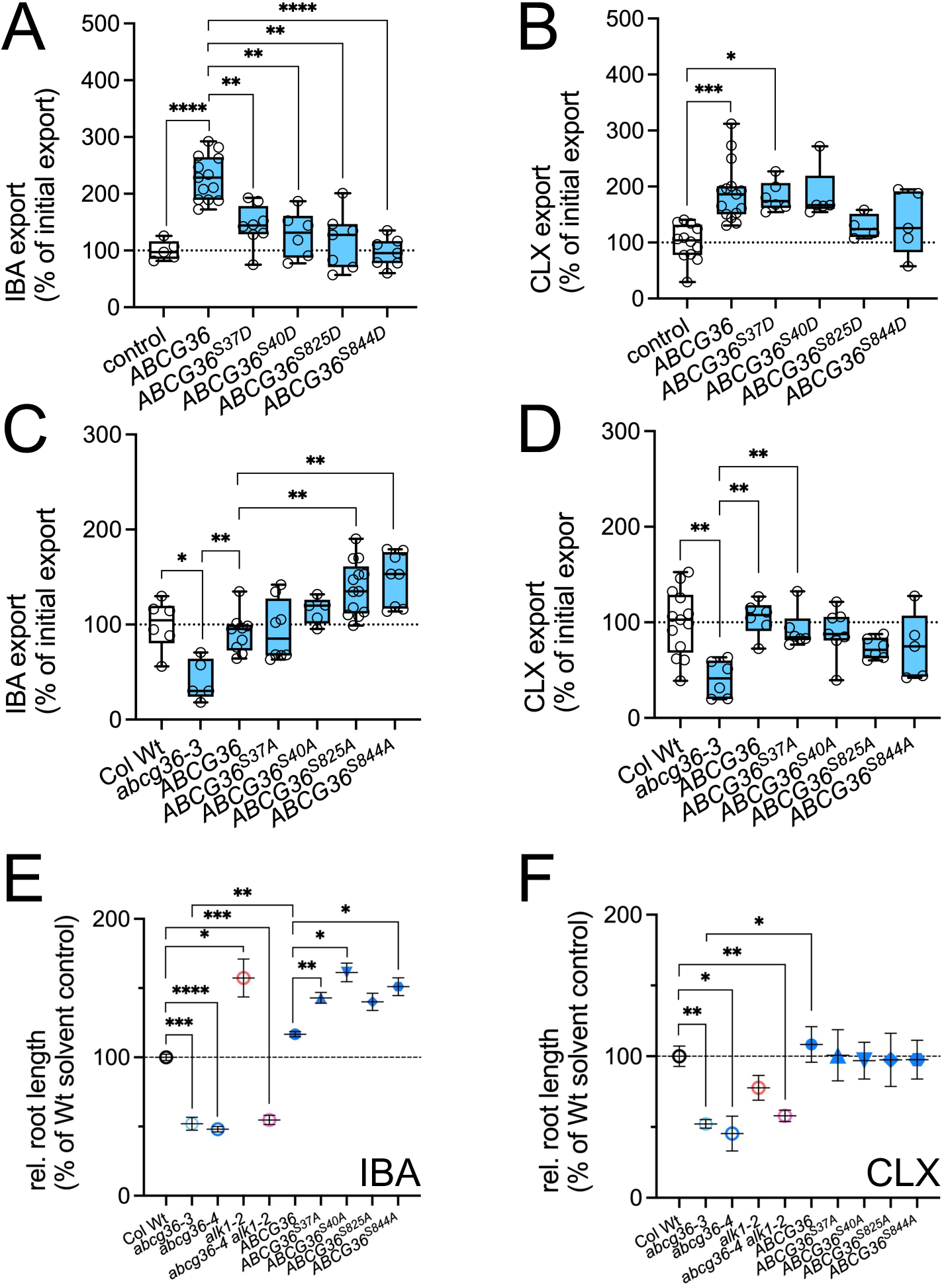
Phosphorylation of ABCG36 by ALK1 inhibits IBA but not CLX transport *in planta*. **(A-B)** IBA (**A**) and CLX (**B**) export from *N. benthamiana* protoplasts after transfection with indicated phospho-mimicry (S-to-D) mutations of *ABCG36* expressed under the constitutive *35SCaMV* promoter. Significant differences (Brown-Forsythe and Welch ANOVA with Dunnet’s T3 multiple comparison test) of mean export ± SE (n ≥ 4 independent transfections) to vector control and Wt ABCG36, respectively, are indicated: *, *p* < 0.05; **, *p* < 0.01; ***, *p* < 0.001; ****, *p* < 0.0001). Controls are in Suppl. Fig. 6. **(C-D)** IBA (**C**) and CLX (**D**) export from *Arabidopsis* protoplasts prepared from indicated phospho-dead (S-to-A) mutations of *ABCG36-GFP (PEN3:PEN3-GFP)* lines in the *abcg36-3* background (Underwood and Somerville, 2017). Significant differences (Brown-Forsythe and Welch ANOVA with Dunnet’s T3 multiple comparison test) of mean export ± SE (n ≥ 4 independent protoplast preparations) to Wt, *abcg36-3* or *ABCG36* in *abcg36-3*, respectively, are indicated: *, *p* < 0.05; **, *p* < 0.01; ***, *p* < 0.001; ****, *p* < 0.0001). Controls are in Suppl. Fig. 6. **(E-F)** Relative root length of indicated Arabidopsis lines grown 12 days on 7.5 μM IBA (**E**) or 5 ug/ml CLX (**F**); Wt growth is set to 100%. Significant differences (unpaired *t* test with Welch’s correction, *p* < 0.05) of mean root lengths ± SE (n ≥ 4 independent transfections) to Wt, *abcg36-3* or *ABCG36* (*PEN3:PEN3-GFP* in *abcg36-3*), respectively, are indicated: *, *p* < 0.05; **, *p* < 0.01; ***, *p* < 0.001; ****, *p* < 0.0001).

Next, we analyzed IBA and CLX export in the *abcg36-3/pen3-3* loss-of-function mutant that was complemented with *ABCG36:ABCG36-GFP* containing the exact S-to-A mutations (Underwood and Somerville, 2017). As expected, Wt *ABCG36:ABCG36-GFP* fully restored mutant export defects for IBA and CLX to Wt level (Fig. 6C, Suppl. Fig. 6C). However, both cluster II but not cluster I phospho-dead mutations provided enhanced IBA (Fig. 6C), while neither cluster I nor cluster II mutations had a significant effect on CLX export that stayed at Wt level (Fig. 6D). Interestingly, combined S44A, S45A and cluster I mutation (containing S37A, S40A, S43A and S45A) showed no difference in IBA or CLX transport in comparison to single cluster I mutations (not shown) indicating the absence of additive effects. As for D mutations, all four phospho-dead mutations did not significantly alter ABCG36 expression or PM location (Suppl. Fig. 6D).

Finally, we aimed at testing the behavior of the S-to-A mutation *in planta* and treated seedlings with cytotoxic (µM) concentrations of IBA and CLX and used root elongation as a read-out for ABCG36 activity detoxifying these substrates (Aryal et al., 2019; He et al., 2019; Lu et al., 2015). In line with enhanced IBA transport rates (Fig. 4), *alk1* provided resistance to IBA in comparison to Wt and hypersensitive *abcg36* alleles and *abcg36 alk1* (Fig. 6E). Again, in agreement with transport data (Fig. 6C), three out of four S-to-A mutations of ABCG36 showed significantly enhanced root growth in comparison to *abcg36-3* complemented with a Wt version *ABCG36* (Fig. 6F). As shown before, both tested *abcg36* loss-of-function alleles were also hypersensitive on CLX (He et al., 2019), while root growth of *alk1* and all four S-to-A mutations of ABCG36 was not significantly different from Wt or complemented knock-out (KO), respectively (Fig. 6F). Overall, this dataset further supports the idea that ABCG36 IBA - but not CLX - transport is regulated by protein phosphorylation requiring ALK1.

### *ABCG36* and *ALK1* loss-of-function but also phospho-dead mutation of *ABCG36* increase *F. oxysporum* susceptibility

Transport and phospho-proteomic data presented so far suggest that ALK1 by means of ABCG36 protein phosphorylation unilaterally shuts off IBA transport allowing for CLX export upon attack. In order to challenge this concept *in vivo*, we soil-infected *abcg36* mutants complemented with phospho-dead versions of *ABCG36* with *F. oxysporum*. As shown previously for *abcg36-4* (Mao et al., 2017a), both *abcg36* loss-of-function alleles revealed significantly increased susceptibility compared to infected Wt indicated by a higher number of disease leaf symptoms, like chlorosis and necrosis (Fig. 7A-B). Interestingly, similar disease sensitivity scores (Suppl. Fig. 7A) were also found for *alk1* but as well for each individual S-to-A mutation of ABCG36 (Fig. 7A-B). Again, S40, S45 and cluster I phospho-dead mutations did not result in additive effects (not shown). Increased disease symptoms correlated well with enhanced vascular penetration of Fo5176 *pSIX1::GFP* in *abcg36* and *ABCG36*^*S40A*^ and *ABCG36*^*S825A*^ lines (Fig. 7C) in comparison to Wt and the hypersensitive, positive control, *agb1-2 (arabidopsis G protein beta 1*; (Trusov et al., 2006)).

**Figure 7:**
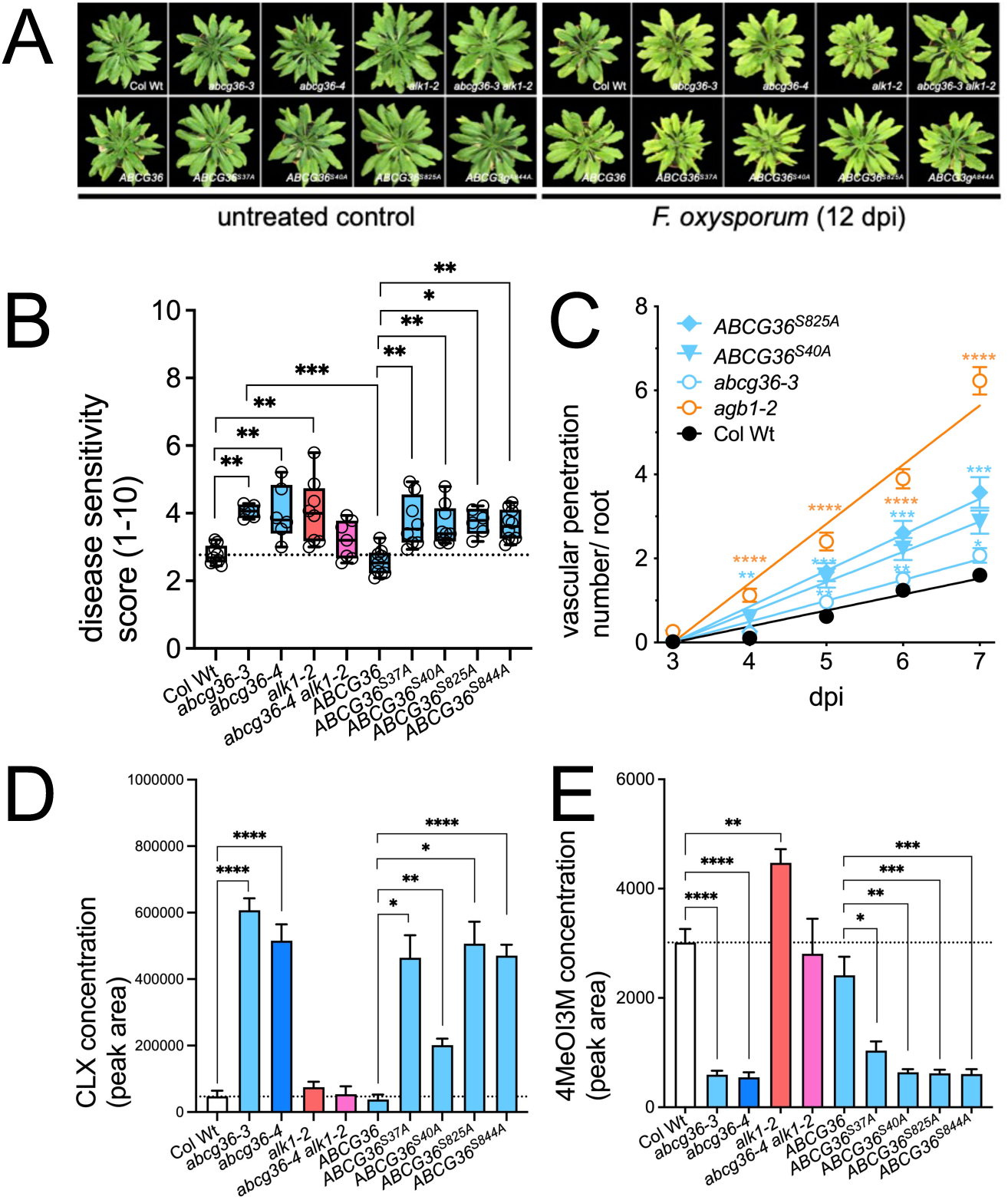
*ABCG36* and *ALK1* loss-of-function - but also phospho-dead mutation of *ABCG36* - increase *F. oxysporum* susceptibility. **(A-B)** 5-week-old plants grown on soil are watered with buffer (untreated control) or with Fo (Fo699 ; 10^7^ conidia/ml). Representative plants are pictured (A) and disease symptoms were quantified using a scale from 0-10 (Suppl. Fig. 7A). Significant differences (Brown-Forsythe and Welch ANOVA with Dunnet’s T3 multiple comparison test) of mean disease symptoms ± SE (n 3 independent infections) to Wt, *abcg36-3* or *ABCG36* in *abcg36-3* are indicated: *, *p* < 0.05; **, *p* < 0.01; ***, *p* < 0.001; ****, *p* < 0.0001). Fresh and dry weight controls are in Suppl. Fig. 7. **(C)** Cumulative root vascular penetration Fo5176 *pSIX1::GFP* in indicated Arabidopsis lines at different days post-transfer to microconidia-containing plates. Significant differences (Ordinary one-way ANOVA) of mean penetration ± SE (n = 3 independent infections) to Wt are indicated: *, *p* < 0.05; **, *p* < 0.01; ***, *p* < 0.001; ****, *p* < 0.0001). **(D-E)** Extracellular levels of camalexin (**D**) and 4MeOI3M (**E**) on *P. infestans*-inocculated *Arabidopsis* leaves determined by non-targeted UPLC-ESI-QTOF-MS. Significant differences (Ordinary one-way ANOVA) of mean levels ± SE (n≥ 7 with each 30 droplets) to wild type, *abcg36-3* or *ABCG36* in *abcg36-3* are indicated: *, *p* < 0.05; **, *p* < 0.01; ***, *p* < 0.001; ****, *p* < 0.0001). Controls are in Suppl. Fig. 8.

Root infection resulted in strongly reduced plant biomass (fresh and dry weight) of *abcg36*, while biomass of *alk1* or phospho-dead mutant alleles was not significantly affected (Suppl. Fig. 7B-C). This is of interest because the difference between disease symptoms and biomass as indicators for defense and growth, respectively, together with enhanced IBA but non-altered CLX transport rates for *alk1* and phospho-dead mutants (Fig. 6), provides strong evidence that phosphorylation of ABCG36 by ALK1 uncouples CLX-mediated defense but not IBA-mediated growth provided by ABCG36 activity. Thus reduced CLX transport is most likely the consequence of elevated IBA transport.

The necrotophic nature of Fo infection did not allow us to verify this apparent unilateral control of substrate specificity on ABCG36 by ALK1 during Fo infection on the transport level. As alternative approach, and in order to quantify the effect of infection on excreted ABC36 substrates *in planta*, we switched back to the *P. infestans* drop-infection system allowing us to quantify excreted substrates by non-targeted metabolomics ((Matern et al., 2019); Suppl. Fig. 8)).

We found that CLX excretion after infection in *alk1* was not significantly different from Wt verifying our transport data under standard conditions (Fig. 4). These data also underline again that CLX export by ABCG36 is not altered by ALK1-mediated phosphorylation. Excitingly, all four individual phospho-dead ABCG36 lines that revealed unaltered CLX transport without infection (Fig. 6D) showed significantly increased CLX excretion in comparison to the complemented KO ((Matern et al., 2019); Fig. 7D)). This might seem counter-intuitive at first, but might be simply due to enhanced IBA transport as discovered in Fig. 6, limiting CLX efflux by competition and therefore triggering the same compensation mechanism as seen in both tested *abcg36* alleles.

Unfortunately, we were unable to detect exported IBA in the infection droplets, most-likely due to its low abundance (Strader and Bartel, 2011). As a second-best option, we looked at the ABCG36 substrates, 4MeOI3M and 4MeOI3Cys, both - based on competition experiments - being most-likely transported in an IBA manner (Matern et al., 2019). Interestingly, both substrates showed an overall inverse excretion pattern in comparison to CLX (Fig. 7E; Suppl. Fig. 8). This pattern is characterized by reduced but enhanced 4MeOI3M excretion in *abcg36* and *alk1* mutants, respectively, in comparison to the Wt and significantly reduced 4MeOI3M found in infection droplets of all four phospho-dead mutants of ABCG36 again in similarity to both *abcg36* alleles (Fig. 7E).

In summary, on one hand, our results clearly indicate a common function of ABCG36 and ALK1 in disease resistance toward the root-penetrating fungus, *F. oxysporum*. On the other they further support the concept that ALK1 shifts ABCG36 substrate specificity in the direction of defense by dampening IBA transport caused by means of ABCG36 protein phosphorylation upon infection.

## Discussion

### ABCG36 is an exporter of multiple indolic compounds, including IBA and camalexin

In analogy to several other Pleiotropic Drug Resistance proteins (PDRs) of the ABCG subfamily, ABCG36 had been suggested to function as a pleiotropic transporter for a series of structurally unrelated substrates, however, transport studies have only been provided for the indolic compounds, IBA (Aryal et al., 2019) and 4MeOI3M (Matern et al., 2019). Here by using cellular tobacco and Arabidopsis as well as *in vitro* transport systems, we provide direct evidence that ABCG36 is a valid exporter of the phytoalexin, camalexin (Fig. 1A, D). These transport data are hardened by CLX-induction of ABCG36 ATPase activity (Fig. 1E) and competition experiments (Suppl. Fig. 3C). Further, CLX strongly upregulates ABCG36 expression (Fig. 3) as found for many ABC transporters (Hwang et al., 2016). However, the ultimate proof for ABCG36 as CLX transporter comes from the finding that both *abcg36-1/pen3-1* and *abcg36-2/pen3-2* alleles containing single point mutations in ABC signature (G345D) and Walker A motif (G915S) of N- and C-terminal NBDs, respectively, essential for ATP binding and hydrolysis (Locher, 2008), are unable to transport CLX (Fig. 1D) and IBA (not shown) despite being present at the PM (Lu et al., 2015).

Recently, it was suggested that ABCG34/PDR6 and ABCG40/PDR12, the latter in conjunction with ABCG36, are themselves exporters of camalexin (He et al., 2019; Khare et al., 2017), although reports were conflicting: while *abcg36 abcg40* double mutants exhibited enhanced susceptibility to the necrotrophic fungus *Botrytis cinerea*, as well as hypersensitivity to exogenous camalexin (He et al., 2019), ABCG34 seemed not to be required for *B. cinerea* resistance and *B. cinerea*–induced camalexin secretion (He et al., 2019). In contrast, ABCG34 was suggested to secrete camalexin from the epidermal cells to the surface of leaves and thereby conferring resistance to *A. brassicicola* infection (Khare et al., 2017). Our analyses indicated that ABCG34 and G40 (as well as ABCG35 and ABCG37) are most likely not CLX exporters *per se* but rather that ABCG34 and ABCG40 might contribute to CLX transport by interfering with ABCG36 transport, thus acting as negative regulators of ABCG36. This is supported by strongly reduced ABCG36-mediated CLX transport upon co-expression with ABCG40 (Fig. 1A) and enhanced CLX export in *abcg40* and *abcg34* (Fig. 1D).

On the other hand, ABCG36 itself could function as a repressor of an unknown CLX exporter that in the absence of ABCG36 (or upon phospho-dead mutation of ABCG36 and upon infection) gets activated. This would explain highly elevated levels of exported CLX in *abcg36* alleles and phospho-dead mutations of ABCG36 upon infection (Fig. 7D; (Matern et al., 2019)). The latter might be recognized in the cell as non-functional CLX exporters (Fig. 4D) leading in analogy to the loss-of-function mutant alleles to an activation of other CLX exporters. The fact that this enhanced CLX export does not result *FOX* resistance (Fig. 7A-B) is most likely due to the fact that this third-party CLX export to the leaf surface might be not directed toward the pathogen or simply indicate root-shoot differences.

Such a regulatory role for ABCGs is new but has been described for another subgroup of mammalian ABCs (Aryal et al., 2015; Principalli et al., 2015). A prominent example is the sulfonylurea receptor (SUR/ABCC8;9) that associates with the potassium channel proteins, Kir6.1 or Kir6.2, to form an ATP-sensitive potassium channel (Principalli et al., 2015). Within the channel complex, SUR/ABCC8;9 serves as a regulatory subunit, which fine-tunes potassium channel gating. For ABCG36, such a regulatory interplay between related ABCG isoforms is supported by iTRAQ analyses of two independent *abcg36* alleles that indicated that ABCG34/PDR6 (but not ABCG37/PDR9) is down-regulated on the protein level (not shown). Interestingly, also ALK1-YFP was shown to pull-down ABCG34 and ABCG40 (Suppl. Table. 2) indicating that several ABCG transporters (together with ALK1) are eventually clustered in a functional complex. Thus, physical and regulatory networks between ABCG transporters might therefore be the reason for a challenging interpretation of the transport data.

### A novel mode of direct ABC transporter substrate specificity regulation by the LRR receptor-like kinase, ALK1/QSK1/KIN7

In search for a regulator that controls activity of ABCG36 towards multiple substrates, we isolated the LRR receptor-like kinase, ALK1/QSK1/KIN7, as an ABCG36 interactor by co-immunoprecipitation and interaction was substantiated by *in planta* co-locations, FRET-FLIM analyses in tobacco and a shared up-regulation by ABCG36 substrates (Figs. 2-3; Suppl. Fig. 1). Co-expression in tobacco and substrate competition experiments uncovered a functional (inhibitory) impact of ALK1 on ABCG36 IBA transport activity that directly depends on ALK1 kinase activity (Fig. 4), which was proven by analyses of IBA transport of *alk1* and phospho-mimicry and phospho-dead mutations of ABCG36 (Fig. 6). Importantly, mutation of phospho-sites on ABCG36 or of ALK1 only affected IBA but not CLX transport, which remained at Wt or vector control level, respectively (Figs. 4, 6). This unilateral regulation of ABCG36-mediated IBA transport by ALK1 phosphorylation was further hardened *in planta* revealing that *alk1* and were unlike *abcg36* resistant to cytotoxic concentrations of IBA but not of CLX (Fig. 6E-F) that based on our transport experiments is most-likely caused by constitutive up-regulation of IBA export. Final proof for our concept in that CLX export is competed out by constitutively enhanced IBA transport comes from *F. oxysporum* root infection analyses that revealed increased disease susceptibility and fungal penetration of the vasculature for *alk1* and phospho-dead *ABCG36* mutant alleles that behaved like *abcg36* alleles (Fig. 7).

We identified two ALK1-specific phosphorylation sites on each of the two phosphor-site clusters preceding the NBDs (Fig. 5) and found that single mutations of all four tested serines on both clusters had both in respect to transport and plant physiology and pathogenicity in principle the same effect (Figs. 6-7). However, it should be stressed that based on our proteomics data in *alk1* (Fig. 5B) and on IBA transport of phospho-dead ABCG36 mutations (Fig. 6C) only cluster II sites (S823 and S825) can be safely attributed as relevant phospho-switches of ABCG36 substrate specificity by ALK1. This is in contrast to a recent study in that amongst the tested ABCG36 phospho-sites of cluster I and II, only S40A and S45A mutations failed to restore *B. graminis* penetration resistance (Underwood and Somerville, 2017). The difference of ABCG36 phosphorylation sites for defense might be caused by the choice of different pathogens and tissues, here the root fungus, *Fusarium*, versus the leaf pathogen powdery mildew (Underwood and Somerville, 2017).

An interesting finding was that cluster I phospho-site mutations neither in transport nor in pathogenicity resulted in additive effects (not shown; (Underwood and Somerville, 2017)) indicating that a single phosphorylation of a serine leads to the same effect (here: inhibition of IBA export) and to a very similar extend, suggesting that ALK1 phosphorylation of ABCG36 has a switch-like function.

Still a puzzling question is how phosphorylation of a single serine on obvious flexible and disordered regions, such as the N-terminus and the linker region of ABCG36 (Suppl. Fig. 4A), can have such a drastic effect on ABCG36 substrate specificity. One explanation might be that these regions function in analogy to the regulatory linker or R-domains connecting the two halves of full-size transporters of ABCBs or CFTR/ABCC7, respectively, and where phosphorylation is thought to add negative charges that by means of conformational changes altering ABC functional capacities (Aryal et al., 2015; Gadsby et al., 2006; Geisler and Hegedus, 2020). This model was very recently supported by cryo-EM imaging of the ABCC, yeast cadmium factor (YCF1), showing that the YCF1 R-domain associates upon phosphorylation with NBD1 altering directly its ATPase activity (Khandelwal et al., 2022). For yeast PDRs/ABCGs, it was shown that NBDs influence transport activity but also substrate specificity (Kolaczkowski et al., 2013). A second plausible line of argumentation can be built around the fact that both phosphorylated clusters form surfaces for protein-protein interactions with regulatory proteins or other transporters as described for the R-domain of CFTR/ABCC7 (Bozoky et al., 2013). The latter might involve even phosphorylation-dependent, functional interaction with ABCG34/PDR6 or ABCG40/PDR12 that might explain conflicting CLX transport data.

Regulation of plant (Christie et al., 2011; Henrichs et al., 2012) and non-plant ABC transporters by protein kinases and thus protein phosphorylation is an evolutionary conserved module (Aryal et al., 2015; Geisler and Hegedus, 2020). However, while for plant members of other transporter classes (such as P-type ATPases, PINs or aquaporins), a regulatory impact by receptor-like kinases has been described (Chen et al., 2021; Isner et al., 2018; Li et al., 2021; Wu et al., 2019), ABC transporter regulation by a receptor-like kinase to our knowledge has not yet been reported. Thus, it seems that such short, direct regulatory circuit between receptor kinases and transporters is a generic modular principle allowing plants to very rapidly adjust to changing environments (Wu et al., 2019). Importantly, while all characterized transporter-RLK circuits have only demonstrated regulatory effects on transport capacities, we here present to our knowledge the first description of altered (ABC) substrate specificity determined by an RLK.

Another noteworthy aspect is that the RLK, ALK1/QSK1/KIN7, based on our and previous findings is implicated by means of protein interaction and phosphorylation in the regulation of diverse transporter types (such as aquaporins, potassium channels and ABCGs) acting in different physiological contexts. This apparent promiscuous behavior might not come as a surprise because ALK1/QSK1/KIN7 is a very abundant protein identified and phosphorylated under a variety of biotic and abiotic stresses (Wu et al., 2019). Therefore, it seems that ALK1/QSK1/KIN7 as co-receptor acts in conjunction with other receptor kinases that are responsible for recruiting different substrates to the respective signaling complexes. This is a typical feature of this kinase family where members are part of complexes in that they are differently involved in sensing and transducing extracellular signals, such as hormones, secreted peptides, and pathogens (Boutrot and Zipfel, 2017; DeFalco and Zipfel, 2021). For example, the well-characterized co-receptor, BAK1/SERK3, is part of several receptor kinase complexes and plays a role in diverse signaling pathways ranging from brassinosteroid signaling to defense or light responses or developmental processes (Boutrot and Zipfel, 2017). Currently, the identity of the receptor in the ALK1/QSK1/KIN7-ABCG36 circuit is unknown. The ALK1/QSK1/KIN7 interactor, SIRK1, is an unlikely candidate as IBA export was not altered in *sirk1* (Fig. 4). However, we have identified several other candidate RLKs as ABCG36 and/or ALK1 interactors (Suppl. Table 1-2) that are currently under investigation.

### The ABCG36- ALK1/QSK1/KIN7 module allows to make growth-defense trade-offs on the molecular level

Initially, the involvement of ABCG36 in both growth and defense programs of Arabidopsis was explained by overlapping functions of exported phytochemicals in plant development and plant defense (Bednarek et al., 2009). In light of recent findings (Aryal et al., 2019; He et al., 2019; Khare et al., 2017) and those presented here, it is more likely that ABCG36 exports multiple substrates, a feature that is not unusual for PDR-type ABCG transporters (Borghi et al., 2015), that are defining its individual role at the crossroad between growth and defense. While the individual role for ABCG36 (Aryal et al., 2019; He et al., 2019; Kadota et al., 2019; Khare et al., 2017; Lu et al., 2015; Matern et al., 2019; Stein et al., 2006; Underwood and Somerville, 2013, 2017) and the general function of LRR-RLKs (DeFalco and Zipfel, 2021; Kadota et al., 2019) in both growth and defense processes is not entirely new, their interplay as a functional module has not yet been demonstrated before. By analyzing the transport and role of two ABCG36 substrates, the growth hormone, IBA, and the phytoalexin, CLX, we demonstrate that the central decision for a plant “to grow or fight” is apparently made on the molecular level by the transporter itself that has the capacity to preferentially select one of the two substrates over the other. This “choice” is interestingly a unilateral one and regulated by ABCG36 phosphorylation provided by the RLK, ALK1/QSK1/KIN7, that functions as a molecular switch for ABCG36 substrate specificity: ABCG36 phosphorylation by ALK1 dampens IBA transport (and thus growth) allowing for CLX transport in the direction of defense. This unilateral mode of action most likely works because based on our transport studies, in an “intermediate phosphorylation” status, IBA seems to be the preferred and thus the “default” ABCG36 substrate (Suppl. Fig. 9B). In a “growth state”, ABCG36 would be dephosphorylated by an unknown phosphatase leading to enhanced IBA transport and thus growth (Suppl. Fig. 9A). Finally, upon attack ALK1 would be activated by a so-far not identified receptor promoting ABCG36 phosphorylation, which blocks IBA transport allowing for CLX export toward the pathogen (Suppl. Fig. 9C).

Interestingly, all four diagnosed phospho-sites, S45 to 825, were strongly dephosphorylated by IBA in comparison to solvent control (Fig. 5B), which again based on our results leads to enhanced IBA export. Contrarily, CLX (though demonstrated only for S45) enhances slightly ABCG36 phosphorylation, which allows for enhanced CLX transport. Thus, it appears that phosphorylation events initiated by both ABCG36 substrates though having inverse effects promote IBA and CLX export, respectively, leading eventually to self-amplifying growth and defense cycles (Suppl. Fig. 9).

Another exciting outcome of our study is that IBA treatment and *Fusarium* infection resulted likewise in strongly reduced ABCG36 phosphorylation (Fig. 5) and ABCG36 up-regulation on the protein level (Fig. 3), suggesting a mechanistic coupling. Reduced ABCG36 phosphorylation enhances IBA but lowers competitively CLX export, which from a plant perspective is counter-intuitive because it weakens fungal defense. Interestingly, Fo elicitor treatment enhanced ABCG36 phosphorylation suggesting eventually that Fo is employing IBA as a tool to shut-down plant defense provided by ABCG36. Support comes from the finding that Fo is known to trigger tissue-specific regulation of auxin transport and signaling genes (Lyons et al., 2015) affecting auxin homeostasis (Kidd et al., 2011). On the other hand, chemical or genetic alteration of polar auxin transport conferred increased resistance to Fo infection, indicating that this fungus requires components of transport to efficiently colonize the plant root (Bitas et al., 2015). In light of these findings, it is tempting to speculate that Fo itself, suggested to produce auxins (including IAA, (Kidd et al., 2011)), manipulates ABCG36 expression by IBA secretion. Although IBA production has to our knowledge not been shown for Fo, it is widely distributed among fungi (Ludwig-Muller, 2015). Finally, such a hijacking strategy of Fo to promote disease development in Arabidopsis has been already shown for jasmonate signaling (Thatcher et al., 2009).

In summary, our findings contribute to our understanding how substrate specificity of ABC transporters is regulated. This is on one hand of interest because many ABC proteins are of clinical relevance because their promiscuous substrate selectivity causes pleiotropic or multidrug resistance phenomena (Ambudkar et al., 2006). On the other hand, the identification of this functional ABCG36-ALK1/QSK1/KIN7 module and its molecular mechanisms standing literally at the crossroad between plant growth and defense might indicate that plants are also forced to balance growth and defense because both molecular components (here: ABCG36 and ALK1/QSK1/KIN7) and not only carbon resources are occupied by both pathways. Such an unusual control of IBA and CLX efflux on the transporter level might be necessary because both indolic ABCG36 substrates are metabolically connected by their common intermediate, IAOx, making a control on the metabolic level difficult. Transfer of this knowledge might be helpful to design new breeding and engineering strategies in order to improve plant fitness (Huot et al., 2014). For centuries, crops have been bred to maximize growth-related traits resulting in a loss of genetic diversity that often compromises defense (Strange and Scott, 2005).

## Material and Methods

### Plant material and growth

The following *Arabidopsis thaliana* lines in ecotype Columbia (Col WT) were used: *abcg36-1/pen3-1* (CS67926; (Stein et al., 2006)), *abcg36-2/pen3-12* (Stein et al., 2006), *abcg36-3/pen3-3* (SALK_110926), *abcg36-4/pen3-4* (SALK_000578), *abcg37-2/pdr9-2* (SALK_050885), *abcg34-2/pdr6-2* (SALK_036087C), *abcg35-1/pdr7-1* (CS66050) and *abcg40-2/pdr12-2* (SALK_005635), *abcg36-4 abcg37-2 (pen3-4 pdr9-2)* (Ruzicka et al., 2010), *abcg36-3 abcg40-2* and *abcg36-3 abcg37-2 abcg40-2* (He et al., 2019), *alk1-2* (GABI-Kat-689A02), *qsk1-1* (SALK_019840), *qsk2-1* (WiscDsLoxHs082_03E), *qsk1-1 qsk2-1* and *sirk1-1* (SALK_125543) (Wu et al., 2019), *35S:ABCG37-GFP* (Ruzicka et al., 2010), *ABCG36:ABCG36-GFP (PEN3:PEN3-GFP)* (Stein et al., 2006), *ABCG36:ABCG36*^*S37A*^*-GFP (PEN3:PEN3*^*S37A*^*-GFP), ABCG36:ABCG36*^*S40A*^*-GFP (PEN3:PEN3*^*S40A*^*-GFP), ABCG36:ABCG36*^*S825A*^*-GFP (PEN3:PEN3* ^*S825A*^*-GFP)* and *ABCG36:ABCG36*^*S844A*^*-GFP (PEN3:PEN3*^*S844A*^*-GFP)* (Underwood and Somerville ; 2017), *AHA2:AHA2-GFP* (Fuglsang et al., 2014), *ABCB1:ABCB1-GFP* (Mravec et al., 2008). For *35S:ALK1-YFP*, ALK1 cDNA (At3g02880) was amplified by using ALK1-s (5′-AT GAA GTA TAA GCG TAA GC-3′) and ALK1-dir-as (5′-TGG AAC GTT CTC TTC TTT CTT TCT C-3′) primers and inserted into the pCR8/GW/TOPO vector, which was recombined into Gateway destination vector pBIN19-YFP. For *35S:ALK1-mcherry*, ALK1 cDNA was amplified using primers ALK1-dir-s (5′-CAC CAT GAA GTA TAA GCG TAA GCT AAG C-3′) and ALK1-dir-as (5′-GTC GGA TAC AGG ATT TGG GGA G-3′) primers. Similarly, for 35S:ALK1^trunc^-mcherry, ALK1 cDNA was amplified by using ALK1-dir-s and ALK1-dir^trunc^-as (5′-TGG AAC GTT CTC TTC TTT CTT TCT C-3′) primers, cloned into the pENTR/D-TOPO vector followed by a recombination into pEG101-mcherry and transformation into *alk1-2* mutant by floral dipping. ABCG34 (At2g36380) and ABCG35 (At1g15210) cDNAs were custom synthesized by GenScript Biotech (Netherlands) in pDONOR207 and recombined into Gateway destination vector pB2GW7 and pK7RWG2-RFP, respectively. *35S:ABCG40-GFP* (At1g15520) was as described in (He et al., 2019).

DR5:GFP previously described in (Ruzicka et al., 2010) was crossed into *abcg36-4 abcg37-2, alk1-2* and *cerk1-2* and isogenic, homozygous lines for the transgene in the F3 generations were used for further analyses. The same strategy was used for crossing of *ABCG36:ABCG36-GFP* into *alk1-2* and *35S:ALK1-mcherry* respectively. *35S:ABCG36 Arabidopsis* lines were constructed by transforming *35S:PDR8* (Kim et al., 2006) into *pen3-4* mutant.

Seedlings were generally grown on vertical plates containing 0.5 Murashige and Skoog media, and 0.75% phytoagar at 16 h (long day) light per day. Developmental parameters, such as primary root lengths and fresh or dry weight, were quantified as described in (Ruzicka et al., 2010). All experiments were performed at least in triplicate with 30 to 40 seedlings per each experiment.

### Fusarium oxysporum growth, elicitor mix preparation and treatment, and root infection assays

*Fusarium oxysporum f. sp. conglutinans* (Fo), strain 699 (Fo699; ATCC 58110) and strain 5176 (Fo5176) was grown as described in (Kesten et al., 2019; Mao et al., 2017a). Fungal elicitor mix was prepared from Fo5176 as described elsewhere (Kesten et al., 2019). *F. oxysporum* infection assays of Arabidopsis roots in soil were performed with strain Fo699 as reported in (Mao et al., 2017a). In short, 3-week-old *Arabidopsis thaliana* plants were infected by pipetting 10 ml conidia suspension (10^7^ conidia/ml) directly into the soil contained in a 125 ml plastic pot harboring a single plant. Fresh-weight and disease sensitivity scores were obtained by measuring rosette weight and by observing chlorotic and necrotic symptoms scored at a scale from 1-10 as number of affected leaves per plant, respectively, two weeks after inoculation. 15-30 plants were employed per genotype per experiments in each of three independent experiments (n = 3).

Vascular penetration assays were performed as described before (Kesten et al., 2019). Briefly, 8 days-old seedlings grown as described above were transplanted to plates with 100 μl of 10^7^ microconidia/ml of Fo5176 pSIX1::GFP. Under a stereomicroscope, we observe the GFP signal from the fungus specifically when it reaches the host vasculature of the main and/or lateral roots with a clear, central, linear pattern. We count each of the individual GFP-labelled events as an individual fungal vascular penetration of the root vasculature

### Transport studies

^3^H-IBA (ARC ART1112, 25 Ci/mmol), ^14^C-IAA (ARC ART0340, 25 Ci/mmol) or ^3^H-camalexin (custom-synthesized by ARC, 56 Ci/mmol) export from *Arabidopsis* and *Nicotiana benthamiana* mesophyll protoplasts was analyzed as described (Henrichs et al., 2012). *N. benthamiana* mesophyll protoplasts were prepared 4 days after agrobacterium-mediated transfection of combinations of indicated plasmids or plasmid combinations. *35S:ABCG36*^*S37A*^ *35S:ABCG36*^*S40A*^, *35S:ABCG36*^*S825A*^ and *35S:ABCG36*^*S844A*^ were custom-synthesized by Twist Bioscience (San Francisco, USA) in pTWIST-Entry and Gateway-recombined into pB2GW7 and pB7FWG2-GFP. Transfections with anti-silencing strain, RK19, were used as vector control. Relative export from protoplasts is calculated from exported radioactivity taken at t = 0, 5, 10, 15 and 20 min. as follows: (radioactivity in the supernatant at time t = x min.) - (radioactivity in the supernatant at time t = 0)) * (100%)/ (radioactivity in the supernatant at t = 0 min.); presented are mean values from >4 independent protoplast preparations at t = 15 min.

IBA and CLX uptake into *Arabidopsis* vesicles prepared from Arabidopsis lines grown as mixotrophic liquid cultures was measured in the absence (solvent) or presence of 1000 x access of nonlabelled IBA and CLX, respectively, as described in (Matern et al., 2019). Root acropetal and basipetal PAT measurements were performed as described (Lewis and Muday, 2009).

### Measurement of ATPase activity

Vanadate-sensitive ATPase activity was measured from microsomes (0.06 mg/ml) prepared from tobacco plants transfected with vector control or *35S:ABCG36* using the colorimetric determination of orthophosphate released from ATP as described previously (HAO2019). The amount of Pi released in the absence (solvent control) or presence of IBA, IAA, CLX or indole (50 μM) was quantified using a Cytation 5 reader (BioTek Instruments).

### Affinity purification of ABCG36-GFP and ALK1-YFP interacting proteins and mass spectrometric analysis

Microsomal fractions were prepared from *ABCG36-GFP* (*PEN3:PEN3-GFP*) roots grown on 1/2 MS medium for 10 days. Roots (∼0.1 g fresh weight) were homogenized with buffer (50 mM HEPES-KOH, pH 7.2, 5 mM EDTA, 400 mM sucrose, protease inhibitor cocktail). The homogenates were centrifuged at 8,000 × *g* at 4°C for 20 min, and the supernatants were centrifuged at 100,000 × *g* at 4°C for 60 min to prepare the microsomal fractions. The pellets were dissolove in 1 mM dithiothreitol with 1 % CHAPS (DOJIN), C12E8 (Sigma), or n-Heptyl-β-D-thioglucopyranoside (Dojin). Co-immunoprecipitation analyses were carried out as described recently using anti-GFP MicroBeads (Miltenyi Biotec, Germany; (Henrichs et al., 2012)) except that bands of interest were size-selected by Flamingo fluorescent gel stain (BioRad) and manually cut out of the gel prior to trypsin digest. LC-MS/MS analyses were performed by using an LTQ-Orbitrap XL-HTC-PAL system. MS/MS spectra were analyzed using the MASCOT server (version 2.2) searching the TAIR8 database (The Arabidopsis Information Resource). The MASCOT search parameters were as follows: set off the threshold at 0.05 in the ion-score cut off, peptide tolerance at 10 ppm, MS/MS tolerance at ± 0.5Da, peptide charge of 2+ or 3+, trypsin as enzyme allowing up to one missed cleavage, carboxymethylation on cysteines as a fixed modification and oxidation on methionine as a variable modification. Proteins were sorted according to their appearance in triple experiments (identified counts) and listed according to their score (Supplementary Table 1).

ALK1-YFP (*35S:ALK1-YFP*) co-IP/LC-MS-MS was carried out identically except that 10 bands were size-selected by silver stain and manually cut out of the gel prior to trypsin digest.

### Protein phosphorylation analyses by LC-MS/MS

For *in-planta* phosphorylation analyses, Columbia Wt or *alk1-2* Arabidopsis was grown in mixotrophic liquid cultures (½ MS, 1% sucrose) for 12 days. In some cases, Wt cultures were treated for 24h with IBA, CLX (each 10 μM) or Fo5176 elicitor (250 ug/ml) for 24h or exposed to Fo699 for 24h by addition of 10^7^ conidia/ml. Total microsomes were prepared and trypsin digest was processed by using the FASP protocol overnight (Wiśniewski et al., 2009); phoshopeptides were enriched by TiO_2_ affinity beads (Hu et al., 2021). The tip flow-through was desalted by STAGE tips for non-phosphopeptide analysis.

Samples were measured by LC-MS/MS using a nanoscale–HPLC on an EASY-nLC 1000 or an EASY-nLC 1200 Liquid Chromatograph connected online to a QExactive (QE) Plus or HF-X mass spectrometer (Thermo Scientific). Peptides were fractionated on a fused silica HPLC-column tip (I.D. 75 μm, New Objective, self-packed with ReproSil-Pur 120 C18-AQ, 1.9 μm (Dr. Maisch) to a length of 20 cm) over a linear gradient from 4-24% ACN in 0.1% formic acid with a flow rate of 250 nl/min within 85 min. All full-scan acquisition was done in the FT-MS part of the MS in the range from m/z 350-1750 with an automatic gain control target value of 10^6^ for QE plus and 3×10^6^ for HF-X and at resolution 70’000 for QE Plus and 120’000 for HF-X. MS acquisition was done in data-dependent mode to sequentially perform MS/MS on the ten (QE plus) or twelve (HF-X) most intense ions in the full scan (Top10/12) in the HCD cell using the following parameters. A normalized collision energy of 25%, AGC target value: 5,000 (QE plus) and 10,000 (HF-X), and a resolution of 17’500 for QE Plus and 30’000 for HF-X. Singly charged and ions with unassigned charge state were excluded from MS/MS. Dynamic exclusion was applied to reject ions from repeated MS/MS selection for 30 s.

All recorded LC-MS/MS raw files were processed together in MaxQuant version 1.4.1.2 with default parameters using the UniProt full-length *Arabidopsis thaliana* database (April 2016), Homo sapiens (March 2016) or the *Mus musculus* database (April 2016). Search parameters were a mass accuracy threshold of 0.5 (MS/MS) and 20 ppm (precursor), Trypsin/P as protease, maximum three missed cleavages, carbamidomethylation (C) as fixed modification and oxidation (M), phosphorylation (STY) and protein N-terminal acetylation as variable modifications. MaxQuant was used to filter the identifications for an FDR below 1% for peptides, sites and proteins using forward-decoy searching. Match between runs were enabled with a retention time window of 2 min. Phosphosites with localization probabilities ≥0.75 (class I sites) were used for bioinformatics analyses. The abundances of phosphopeptides were normalized to protein level.

### ALK1/QSK1 and SIRK1 in vitro protein phosphorylation analyses

*His-GST-SIRK1* and *His-GST-QSK1* (Wu et al., 2019) were transformed into *E. coli* BL21 (DE3), induced for 3 h induction by IPTG (isopropyl b-D-thiogalactopyranoside) and soluble extracts were purified using GST-sepharose 4B (GE Healthcare, Sweden) as described elsewhere (Geisler et al., 2003) Kinase activities were detected by kinase assays on peptide level. Purified His-GST-SIRK1 and His-GST-QSK1 were resuspended in 100 ul kinase reaction buffer with ATP (50 mM Tris/HCl pH 7.5, 10 mM MgCl2, 1 mM MnCl2 2 mM DTT, 100 μM ATP and 1 x PhosSTOP (Roche, Germany) and as kinase substrates, 20 µl of mixture of 1 fM of each synthetic peptide or peptides from trypsin digested MEFs whole proteome or lamba phosphatase treated peptides from trypsin digested HeLa whole proteome was added. ATP was omitted from control samples. Kinase assays were performed at 30° C for 1 h. Reactions were quenched by addition of 15 µl 10% TFA and spun at 21’000 g for 5 min. Peptides in the supernatants were desalted and purified by C18 (3M™ Empore™, US) packed tips. Peptides were eluted from the tips using 0.1% TFA in 80% ACN. Phosphopeptides were enriched by TiO_2_ affinity beads for LC-MS/MS analysis.

For the analysis of synthetic peptides, the selected-ion monitoring chromatogram (SIM) of each phosphorylated peptide were extracted and compared to control samples (-ATP). The m/z of MS2 were matched to UCSF MS-Product (https://prospector.ucsf.edu/prospector/cgi-bin/msform.cgi?form=msproduct) generated fragments. For the peptide library analysis, the intensities of phosphorylated peptides were compared to control samples (-kinase). The sequences of peptides, which had more than 4-fold changes were chosen for sequence logo analysis using PhosphoSitePlus (https://www.phosphosite.org/sequenceLogoAction.action).

### Drop-inoculation of A. thaliana leaves with zoospore suspensions of P. infestans

Rosette leaves of 5-week-old *A. thaliana* plants were drop-inoculated with a zoospore suspension of *P. infestans* Cra208m2 (Si-Ammour et al., 2003) by applying 10 ul drops (5 × 10^5^ spores ml^-1^) onto the adaxial leaf surface. Thirty drops each were collected and immediately frozen in liquid nitrogen for storage. For extraction, the drops were evaporated to dryness and samples were dissolved in 60 ul of 30% methanol, sonicated for 15 min. and analyzed by LC-MS.

### Metabolite profiling using LC-MS

Untargeted metabolite profiling was performed as described (Matern et al., 2019). Targeted relative quantification of metabolites was done on the peak areas of characteristic extracted ion chromatograms in Bruker’s QuantAnalysis V4.1 software.

### Analyses of auxin responses

Homozygous generations of *Arabidopsis abcg36-4, abcg36-4 abcg37-2, alk1-2* expressing DR5:GFP were obtained by crossing with DR5:GFP lines (Ottenschlager et al., 2003). Seedlings were grown vertically for 5 dag and for 4h on 5 μM IBA plates and analyzed by confocal laser-scanning microscopy.

### Confocal laser scanning and FRET-FLIM lifetime imaging

For imaging, seedlings were generally grown for 5dag on vertical plates containing 0.5 Murashige and Skoog media, 1% sucrose, and 0.75% phytoagar at 16h (long day) light per day. For chemical treatments, seedlings were transferred for 12h on test plates containing the indicated chemicals or the solvents. For confocal laser scanning microscopy work, an SP5 confocal laser microscope was used. Confocal settings were set to record the emission of GFP (excitation 488 nm, emission 500– 550 nm), YFP (excitation 514 nm, emission 524–550 nm), mCherry (excitation 587 nm, emission 550–650nm), RFP (excitation 561 nm, emission 600-680 nm). and FM4-64 (excitation 543 nm, emission 580–640 nm).

Distribution of polar ABCG36-GFP expression in the root was quantified with a self-written plug-in by creating a mask for whole roots using the fluorescence image by local contrast enhancement (CLAHE), followed by setting auto threshold (Huang method) to segment the root and perform a binary morphological operation. This step is followed by constructing inner and outer region of interest using distance ratio and finally measuring signal ratio in Fiji REF.

For FRET analysis, binary vectors *ABCG36:ABCG36-GFP, 35S:ALK1-mCherry* and *AHA2:AHA2-RFP* and *p19* as gene-silencing suppressor were transformed into *Agrobacterium tumefaciens* strain GV3101 and infiltrated into *Nicotiana benthamiana* leaves. The measurements were performed 3 dai using a SP8 laser scanning microscope (Leica Microsystems) with LAS-AF and SymPhoTime software as described (Veerabagu et al., 2012). Before FRET-FLIM measurement, the presence of the fluorophores was detected using 488- or 561-nm lasers for GFP or RFP excitation, respectively. The lifetime τ [ns] of either the donor only expressing cells or the donor-acceptor pairs was measured with a pulsed laser as an excitation light source of 473 nm and a repetition rate of 40 MHz (PicoQuant Sepia multichannel picosecond diode laser, PicoQuant Picoharp 300 TCSPC module, and Picosecond event timer). The acquisition was performed until 1000 photons in the brightest pixel were reached. To obtain the GFP fluorescence lifetime, data processing was performed with SymPhoTime software and mono-exponential curve fitting, correction for the instrument response function, and a fitting range from channel 90 to 1400.

### Real-time PCR analyses

Total RNA from plants infected with *F. oxysporum* were isolated by using TRIZOL reagent (Roth, Germany). First strand cDNA synthesis from DNAse-treated RNA (New England Biolabs) was performed with the Omniscript RT kit (Qiagen, Hilden Germany) according to the manufacturer guidelines. qPCR reaction was operated by using the SensiMix SYBR® from Bioline (www.bioline.com) and a Magnetic Induction Cycler (Bio Molecular Systems) using primers listed in Table S2. The PCR condition included initial denaturation cycle for 10 min at 95°C followed by 45 cycles of denaturation for 20s, annealing for 20s at 59°C, extension for 20s at 72°C. Data were analyzed with the software micPCR vers. 1.4.0. Expression of target genes were normalized to the *RHIP1* reference gene ((Czechowski et al., 2005). Three biological replicates (with three technical replicates for each experiment) were performed for each time point after infection.

### Data Analysis

Data were statistically analyzed using Prism 9.3.1 (GraphPad Software, San Diego, CA).

## Acknowledgments

We would like thank L. Charrier (UNIFR) and U. Smolka (IPB) for excellent technical assistance, B. Egger and F. Meyenhofer (both at UNIFR) for help with confocal imaging and quantification of root GFP signals, Silke Lehmann and Jean-Pierre Métraux (both at UNIFR) for initial help with establishing *Fusarium* cultures, M. Fujiwara (Nara Institute of Science and Technology) for help with co-IPs, J. Schneuwly, E. Juttin, L. Affolter, N. Simonet (UNIFR) and T. Romeis (IPB) for experimental support, A. Fuglsang University of Copenhagen) for *AHA2:AHA2-GFP* lines, J. Friml (IST Austria) for *35S:ABCG37-GFP, abcg36-s/pdr9-2, abcg36-4/pen3-4, and abcg37-2/pdr9-2* lines, B. Underwood (University of California) for *PEN3:PEN3-GFP* plasmid and *PEN3* serine mutant lines, W. Schulze (University of Hohenheim) for ALK1/QSK1 and SIRK1 expression plasmids, X. Meng (Shanghai Normal University) for *abcg* double and triple mutant lines and M. Hothorn (University of Geneva) for mutamt consulting. This work was supported by grants from the Swiss National Funds (project 31003A_165877 and 310030_197563 to MG and project 310030_184769 to CSR), the *Pool de Recherche* of the University of Fribourg (to MG), the German Research Foundation (grant RO 1172/6−1 to SR and grants CRC 1101/D02, HA 2146/22, INST 37/819-1 FUGG, INST 37/965-1 FUGG to KH), the Polish National Science Centre (grant 2017/27/B/NZ1/01090 and 2021/41/N/NZ1/04030 (PRELUDIUM 20) to MJ), the Polish National Center for Research and Development (grant POWR.03.02.00-00-1032/16 to KP) and the ETHZ foundation (grant 0-828 20172-16 to CSR).

## Author contribution

MG, BA and JX designed research; BA, JX, ZH, JL, JH, YF, NG, HYH, GSA, KP, MZ, KG and MG performed research; AB, JX, ZH, JL, JH, JF, NG, HYH, GSA, KP, MZ, SR and MG analyzed data; TN, KH, CSR, MJ, SR and MG supervised work; MH designed kinase mutations, MG wrote the manuscript, all authors commented on the manuscript.

## Abbreviations

IBA,: indole-3-butyric acid;
IAA,: indole-3-acetic acid;
PAT,: polar auxin transport;
ABCG,: ATP-binding cassette protein subfamily G;
PDR,: Pleiotropic drug resistance;
ABCB,: ATP-binding cassette protein subfamily B;
PIN,: pin-formed; dag, day(s) after germination;
dai,: day(s) after infection;
WT,: wild-type;
MS,: mass spectrometry.

## Figure Legends

**The following Supporting Information is available for this article:**

**Supplementary Figure 1: ABCG36 and ALK1 co-localize and interact but ALK1 does not alter ABCG36-GFP expression and location**.

**Supplementary Figure 2: *Fusarium oxysporum* root infection alters ABCG36 expression and polarity on the posttranscriptional level**.

**Supplementary Figure 3: IAA export in LRR-RLK loss-of-function mutants and auxin responses in *alk1***.

**Supplementary Figure 4: ALK1/QSK1/KIN7 is an active protein kinase**

**Supplementary Figure 5: ALK1/QSK1/KIN7-specific phosphorylation sites on ABCG36**.

**Supplementary Figure 6: Phosphorylation of ABCG36 by ALK1 inhibits IBA but not CLX transport *in planta***.

**Supplementary Figure 7: Disease symptom scale and fresh and dry weight controls of *F. oxysporum* infections**.

**Supplementary Figure 8: Extracellular levels of metabolites on *P. infestans-*inocculated Arabidopsis leaves**.

**Supplementary Figure 9: Hypothetical model on the action of ALK1/QSK1/KIN7 on ABCG36 phosphorylation**

**Supplementary Table 1: ABCB36/PEN3/PDR8-interacting proteins identified by co-immunoprecipitation using ABCG36-GFP (PEN3:PEN3-GFP) lines followed by LC-MS/MS**.

**Supplementary Table 2: ALK1-interacting proteins identified by co-immunoprecipitation using ALK1-YFP as a bait followed by LC-MS/MS analyses**.

**Supplementary Table 3: Quality control of phospho-proteomics**.

**Supplementary Table 4: Forward (fw) and reverse (rev) primers used for real-time PCR analyses in this study (see Suppl. Fig. 2B)**.

