## Supplemental Information for "A phospho-switch provided by LRR receptor-like kinase, ALK1/QSK1/KIN7, prioritizes ABCG36/PEN3/PDR8 transport toward defense"

### Supplementary Figures

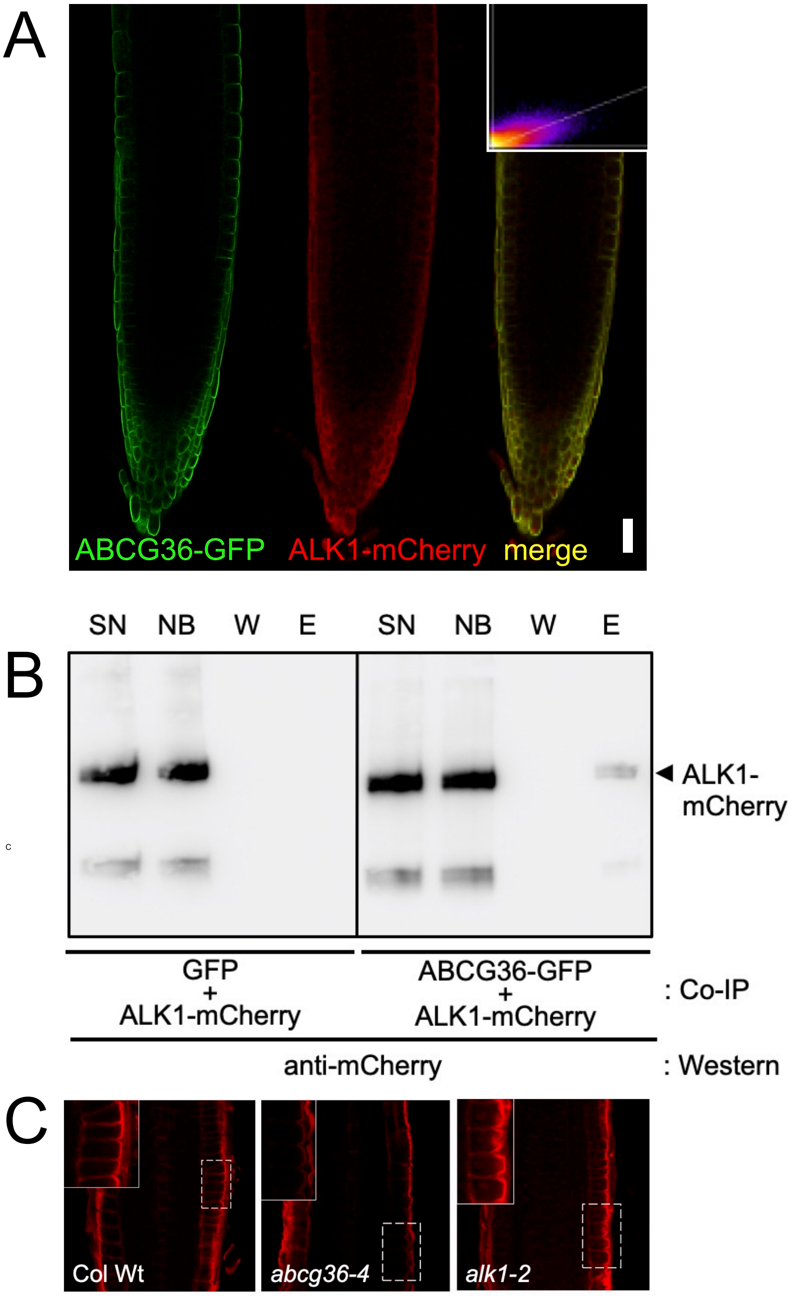


**Supplementary Figure 1: ABCG36 and ALK1 co-localize and interact but ALK1 does not alter ABCG36-GFP expression and location.**
**(A)** Confocal imaging of ABCG36-GFP (*PEN3:PEN3-GFP*) and ALK1-mCherry (*35S:ALK1-mCherry*) stably expressed in Arabidopsis; bar, 100 µm.
**(B)** ABCG36-GFP (*PEN3:PEN3-GFP*) but not GFP alone is able to pull-down ALK1-mCherry (*35S:ALK1-mCherry*) after co-transfection in tobacco (*N. benthamiana*) leaves.
**(C)** ABCG36 expression and localization is not significantly altered in seedling of *alk1-2* comp ared to the Wt (Col Wt) based on immunolocalization of ABCG36 using anti-PDR8 (Agrisera). Note absence of PM signal in *abcg36-4.*

A

| | GFP intensity mock | GFP intensity Fo |
|---|---|---|
| left epidermis | 21.05 | 77.89 |
| stele | 4.66 | 42.59 |
| right epidermis | 20.85 | 60.72 |
| mean (left + right epidermis) | 20.95 | 69.31 |
| epidermis/ stele (ratio) | 4.5 | 1.63 |

B

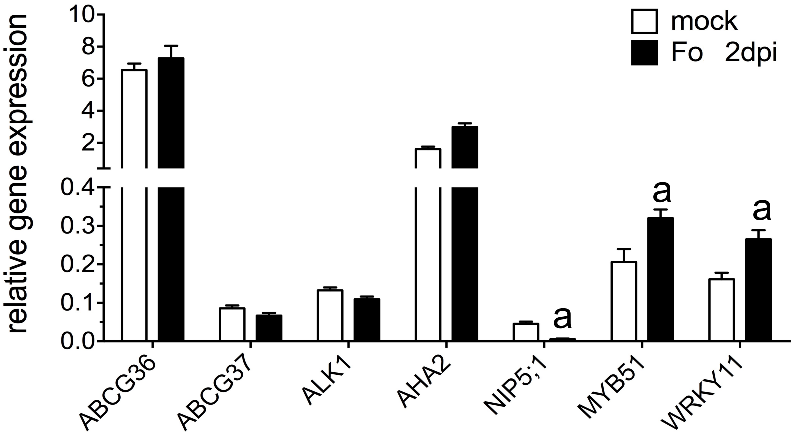


**Supplementary Figure 2: *Fusarium oxysporum* root infection alters ABCG36 expression and polarity on the posttranscriptional level.**
**(A)** Quantification of ABCG36-GFP (*PEN3:PEN3-GFP*) expression and polarity in the Arabidopsis epidermis upon *Fusarium* (Fo) infection.
**(B)** Relative ABCG36, ALK1 and marker gene expression quantified by Q-PCR in Col Wt plants infected with *Fusarium* (Fo). Expression of target genes were normalized to the *RHIP1* reference gene (Czechowski et al., 2005). Three biological replicates (with three technical replicates for each experiment) were performed for each time point after infection. Significant differences of mean ± SE (n =3 RNA preparations) between Fo infection and water control (mock) are indicated by an “a” (unpaired *t* test with Welch’s correction, $p < 0.05$).

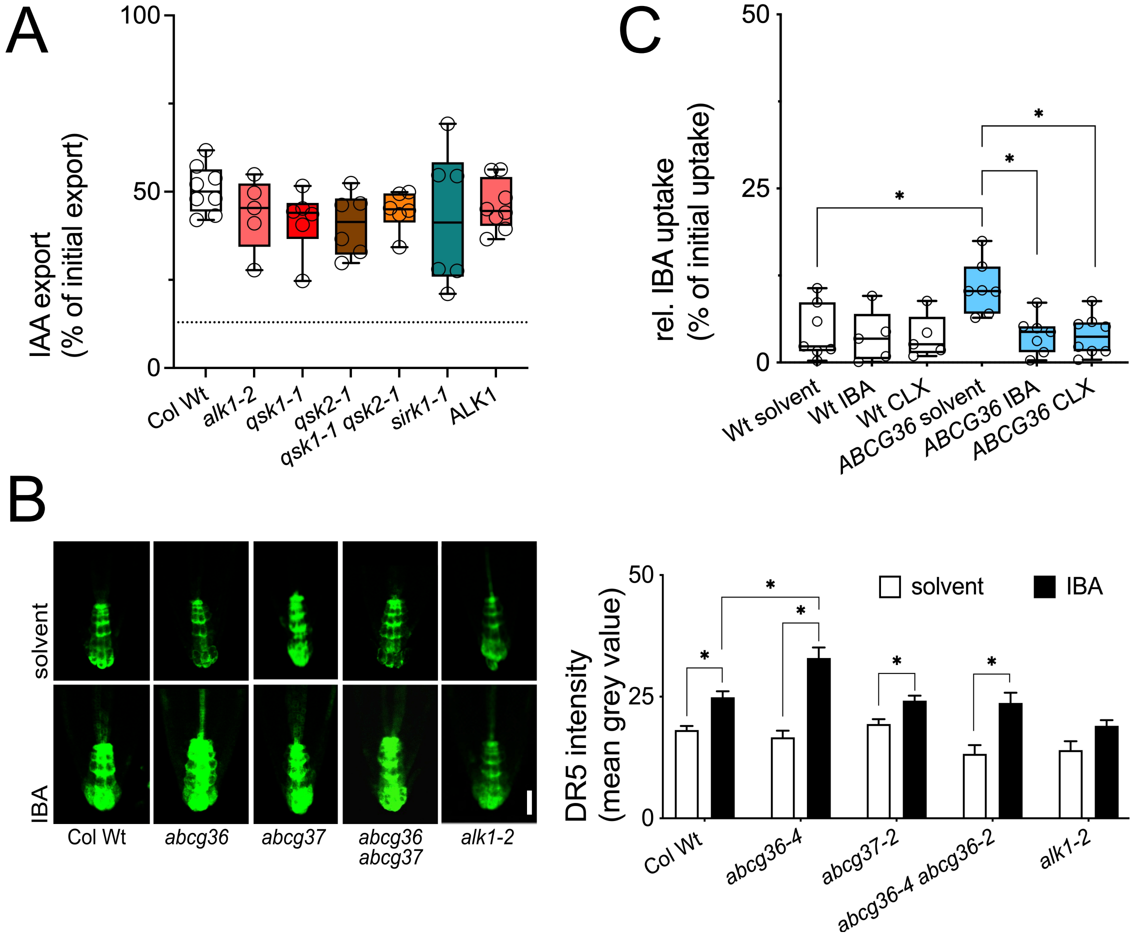


**Supplementary Figure 3: IAA export in LRR-RLK loss-of-function mutants, auxin responses in *alk1* and competition of IBA transport**

**(A)** IAA export from Arabidopsis protoplasts prepared from indicated *LRR-RLK* loss-of function alleles. Differences (Ordinary one-way ANOVA) of mean export ± SE (n ≥ 4 independent protoplast preparations) to wild-type (Col Wt) are non-significant: $p > 0.05$).

**(B)** Root auxin responses visualized (left panel) and quantified (right panel) by the auxin-responsive element, *DR5:GFP*, in the columella of 5 dag seedlings grown on solvent (DMSO) or 5 µM IBA. Significant differences (Ordinary one-way ANOVA) of mean levels ± SE (n = 3 with each > 15 infections) are indicated: *, $p < 0.05$; **, $p < 0.01$; ***, $p < 0.001$; ****, $p < 0.0001$); bar, 20 µm.

**(C)** Competition assays of IBA uptake into *Arabidopsis* vesicles. Uptake of radiolabeled IBA into vesicles prepared from Wt or *35S:ABCG36* lines was measured in the absence (solvent) or presence of 1000 x access of nonlabelled IBA and CLX. Significant differences (Ordinary one-way ANOVA) of mean uptake ± SE (n ≥ 4 independent transfections) to Wt or *35S:ABCG36* solvent control, respectively, are indicated: *, $p < 0.05$; **, $p < 0.01$; ***, $p < 0.001$; ****, $p < 0.0001$).

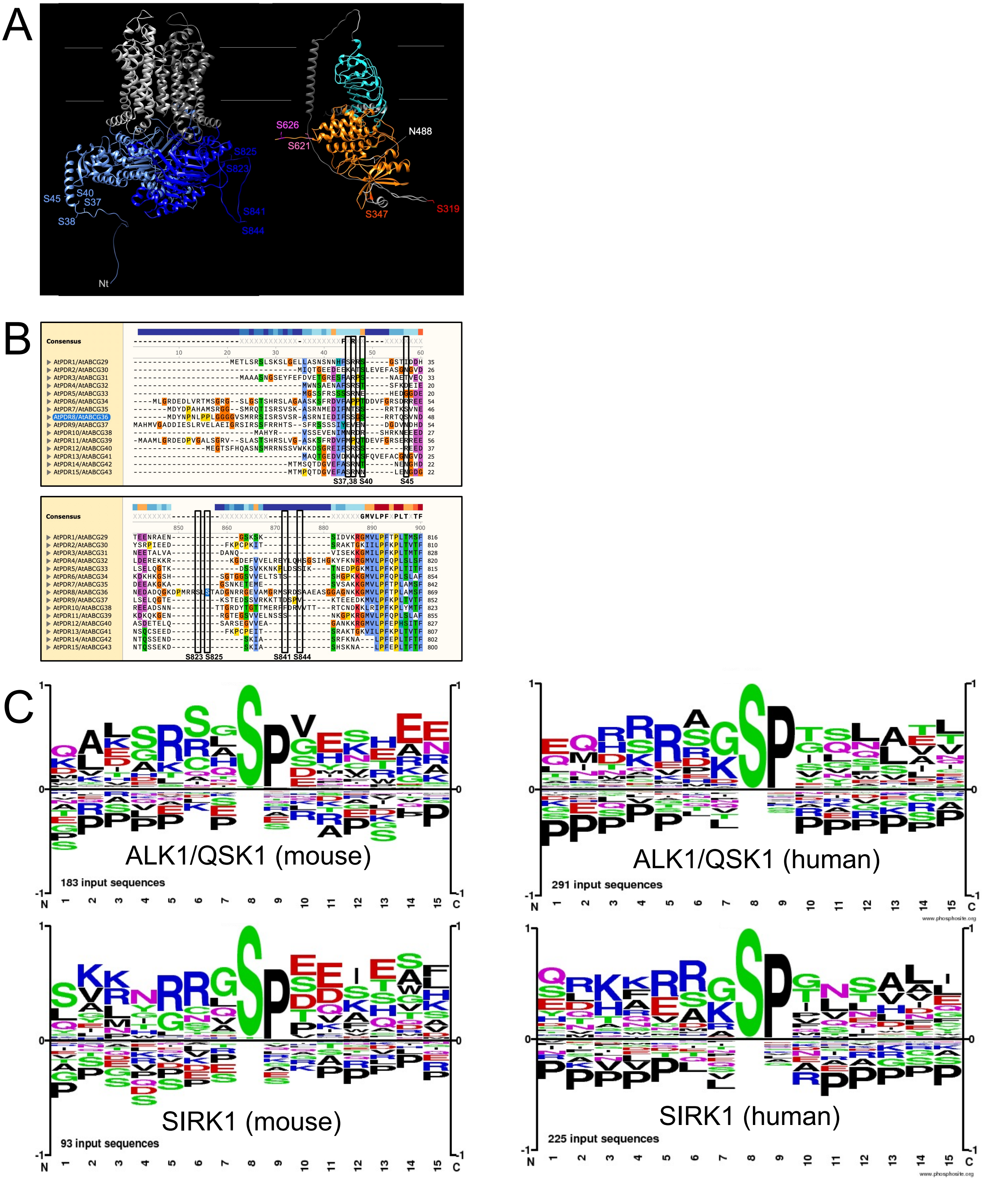


**Supplementary Figure 4: ALK1/QSK1/KIN7 is an active protein kinase**

**(A)** Alpha Fold2 models of ABCG36 (left) and ALK1/QSK1/KIN7 (right) with relevant phospho-sites. Note that the location of apoplastic ecto-domain of ALK1/QSK1/KIN7 is not correctly modeled by Alpha Fold2.

**(B)** Multiple sequence alignment (MegAlign) of cluster I and cluster II phospho-sites in all Arabidopsis full-size ABCGs.

**(C)** *In vitro* kinase activity assays using purified His-GST-SIRK1 and His-GST-ALK1/QSK1/KIN7 and a mouse and a human peptide library, respectively; -kinase was used as a negative control. Protein phosphorylation was analyzed by LC-MS/MS and ALK1/QSK1/KIN7 and SIRK1-specific phosphorylation sites were visualized using the motif analysis tool of phosphosite.org.

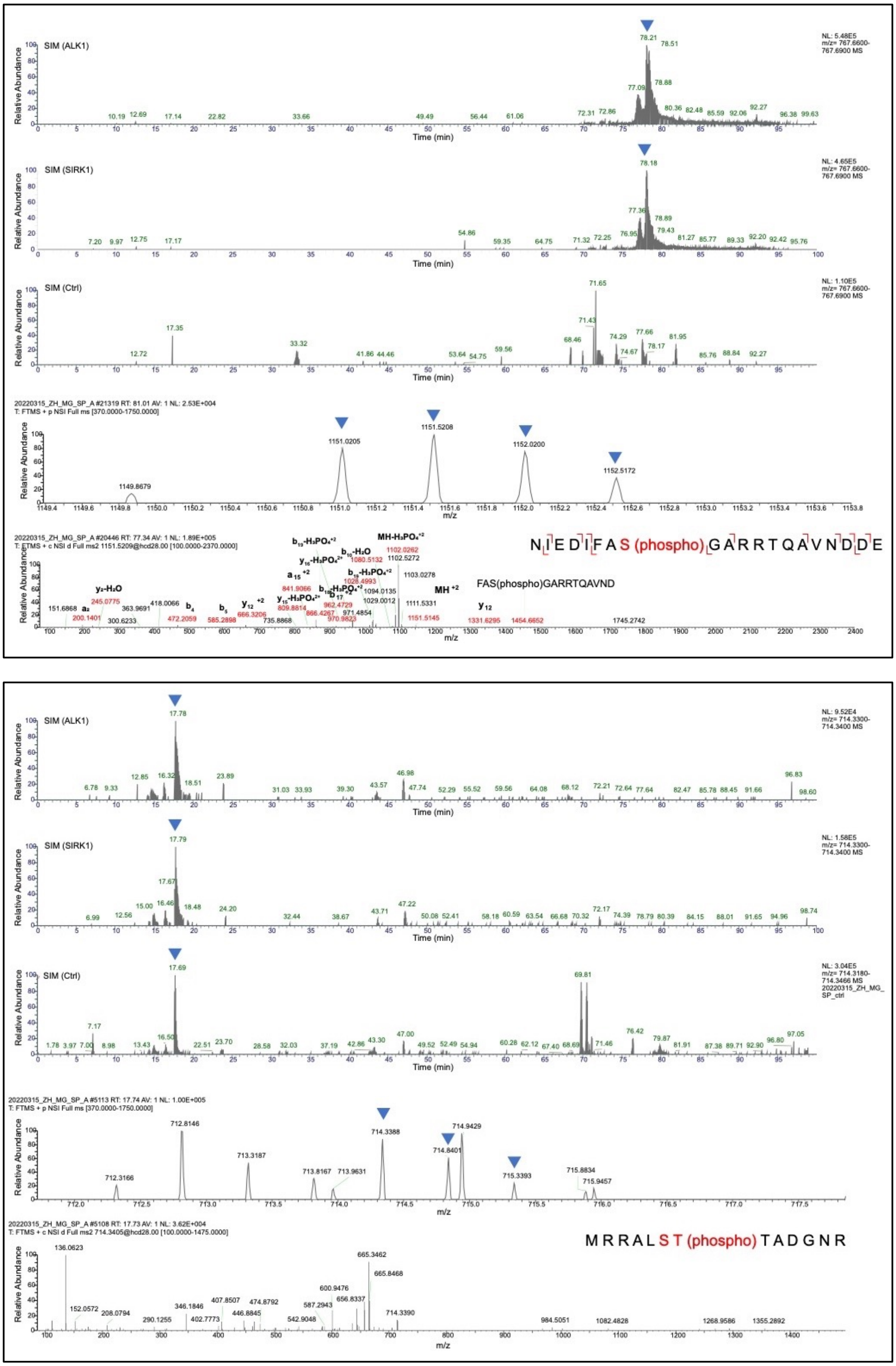


**Supplementary Figure 5: ALK1/QSK1/KIN7-specific phosphorylation of ABCG36.** **(A-B)** *In vitro* kinase activity assays using purified His-GST-ALK1/QSK1/KIN7 and His-GST-SIRK1 with indicated synthetic peptides covering cluster I **(A)** and cluster II **(B)** phosphorylation sites, respectively. ALK1/QSK1/KIN7 and SIRK1-specific phosphorylation sites were analyzed by LC-MS/MS; -ATP was used as a negative control (Ctrl). Shown are selected-ion monitoring (SIM) chromatograms of each phosphorylated peptide compared to control samples (-ATP). S37, S40 and S45 of the cluster I peptide and S823 of the cluster II peptide were mutated on purpose to Ala in order to allow for a safe assignment of phosphorylation sites. Note that the peak corresponding to S38 (blue arrow) is not found in the –ATPase control, while the peak corresponding to S825 (blue arrows) is partially also found in the control.

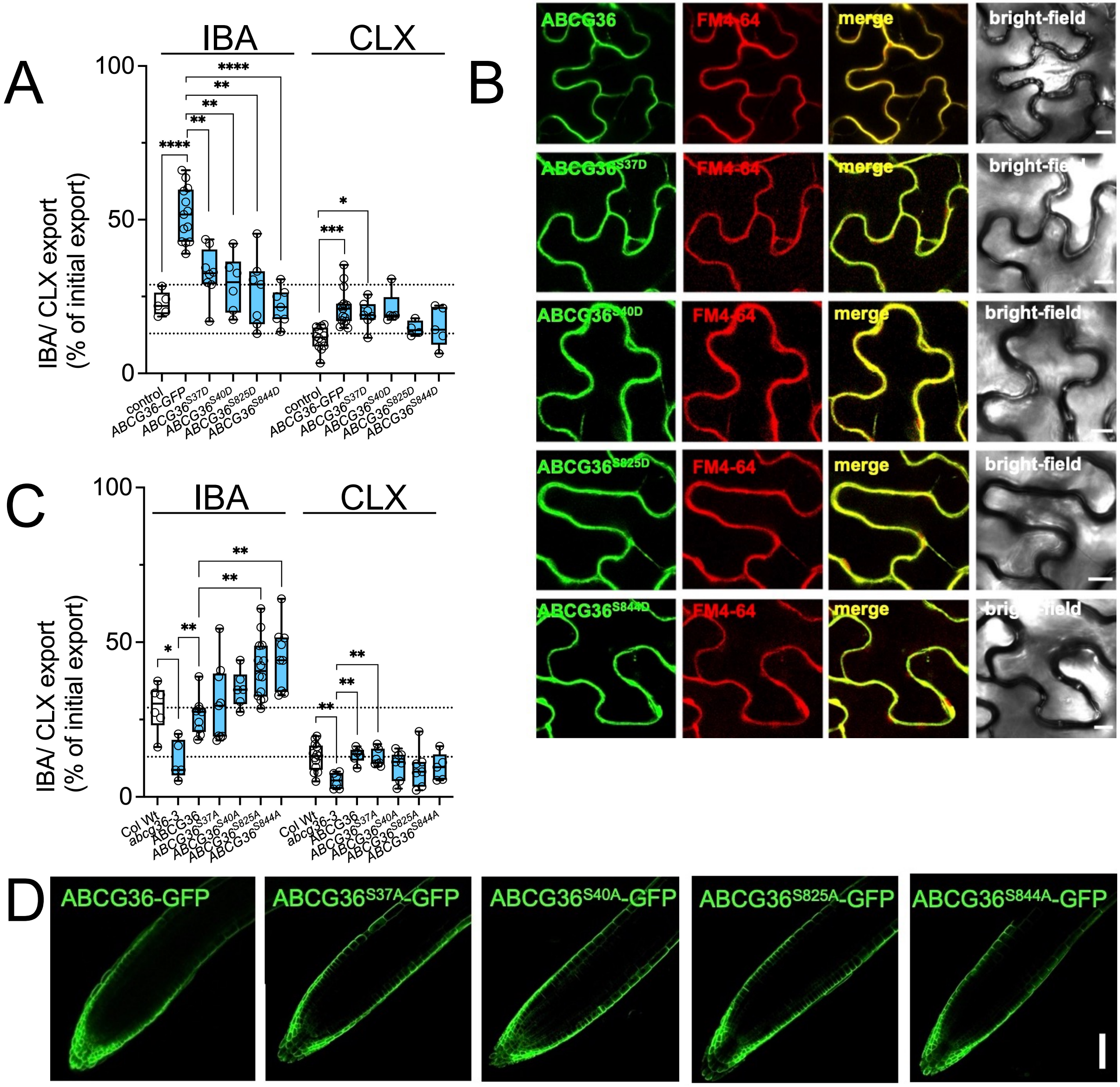


**Supplementary Figure 6: Phosphorylation of ABCG36 by ALK1 inhibits IBA but not CLX transport *in planta***

**(A-B)** Absolute IBA and CL export (**A**) from *N. benthamiana* protoplasts after transfection with indicated phospho-mimicry (S-to-D) mutations of *ABCG36* expressed under the constitutive *35SCaMV* promoter. Significant differences (Brown-Forsythe and Welch ANOVA with Dunnet's T3 multiple comparison test) of mean export ± SE (n ≥ 4 independent transfections) to vector control and Wt *ABCG36*, respectively, are indicated: *, $p < 0.05$; **, $p < 0.01$; ***, $p < 0.001$; ****, $p < 0.0001$). Phospho-mimicry (S-to-D) mutation does not significantly alter plasma membrane expression and location of ABCG36 (**B**); bar, 50 µm.

**(C-D)** Absolute IBA and CLX export (**C**) from Arabidopsis protoplasts prepared from indicated phospho-dead (S-to-A) mutations of ABCG36-GFP (*PEN3:PEN3-GFP*) lines in the *abcg36-3* background. Significant differences (Brown-Forsythe and Welch ANOVA with Dunnet's T3 multiple comparison test) of mean export ± SE (n ≥ 4 independent protoplast preparations) to Wt, *abcg36-3* or *ABCG36* in *abcg36-3*, respectively, are indicated: *, $p < 0.05$; **, $p < 0.01$; ***, $p < 0.001$; ****, $p < 0.0001$). Phospho-dead (S-to-A) mutations does not significantly alter plasma membrane expression and location of ABCG36 (**D**); bar, 200 µm.

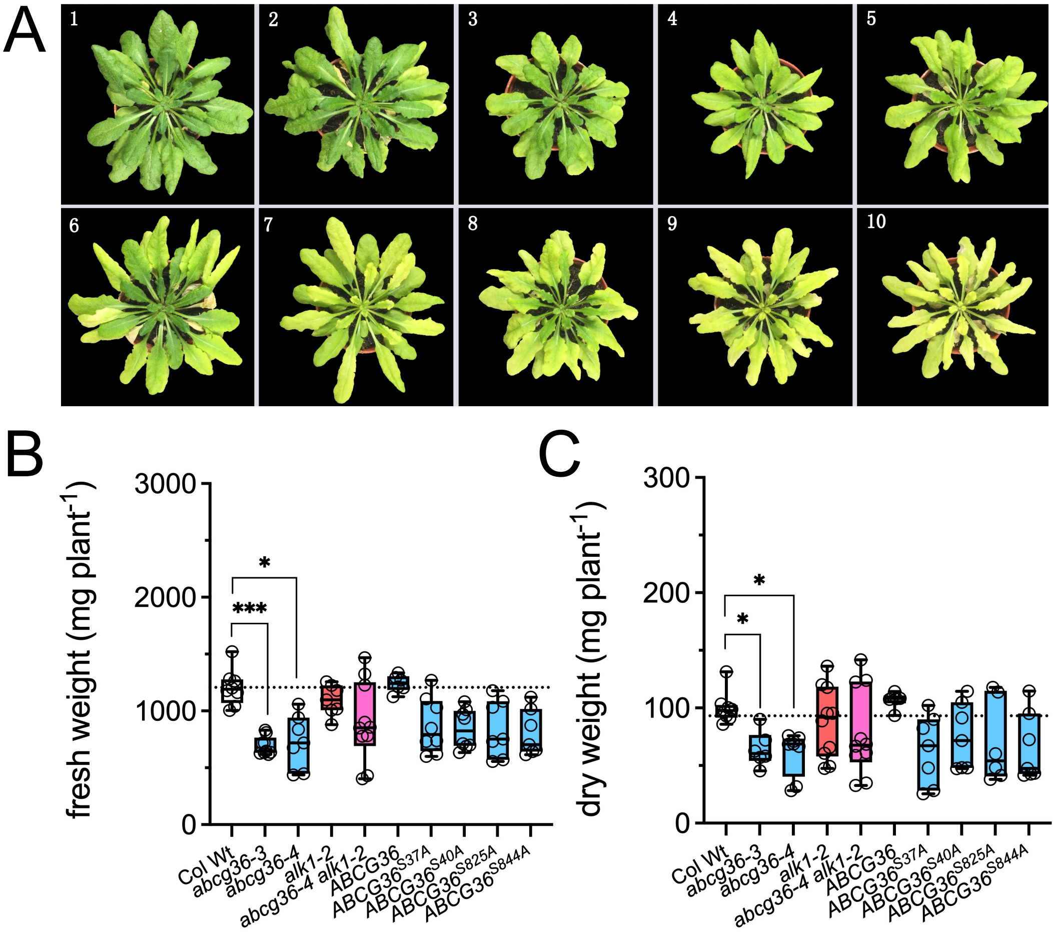


**Supplementary Figure 7: Disease symptom scale and fresh and dry weight controls of *F. oxysporum* infections**

**(A)** 5-week-old Col Wt plants grown on soil are watered with with *F. oxysporum* (Fo699; $10^7$ conidia/ml) and and leaf disease symptoms were evaluated are grouped into categories 1-10.

**(B-C)** 5-week-old plants grown on soil are watered with buffer (untreated control) or with *F. oxysporum* (Fo699; $10^7$ conidia/ml) and fresh (**B**) and dry weight (**C**) were quantified. Significant differences (Brown-Forsythe and Welch ANOVA with Dunnet's T3 multiple comparison test) of mean disease symptoms ± SE (n 3 independent infections) to Wt, *abcg36-3* or *ABCG36* in *abcg36-3* are indicated: *, $p$ < 0.05; **, p < 0.01; ***, $p$ < 0.001; ****, $p$ < 0.0001).

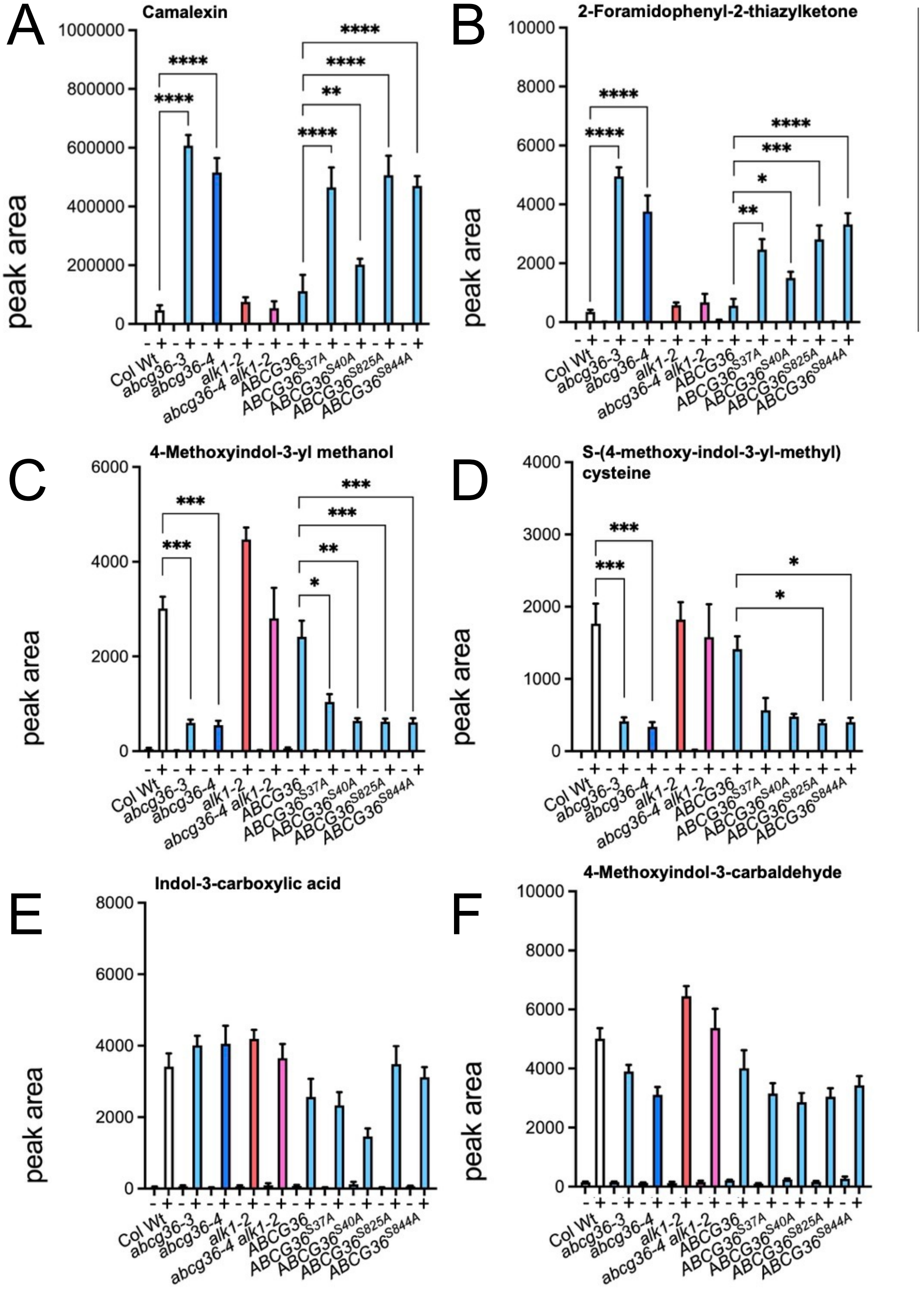


**Supplementary Figure 8: Extracellular levels of metabolites on *P. infestans*-inocculated Arabidopsis leaves.**

(**A-F**) Extracellular levels of indicated metabolites on control (**-**) and *P. infestans*-inocculated Arabidopsis leaves (**+**) determined by non-targeted UPLC-ESI-QTOF-MS. Significant differences (Ordinary one-way ANOVA) of mean levels ± SE (n = 3 with each > 15 infections) to Wt, abcg36-3 or ABCG36 in abcg36-3 are indicated:*, p < 0.05; **, p < 0.01; ***, p < 0.001; ****, p < 0.0001). Controls are in Suppl. Fig. 8.

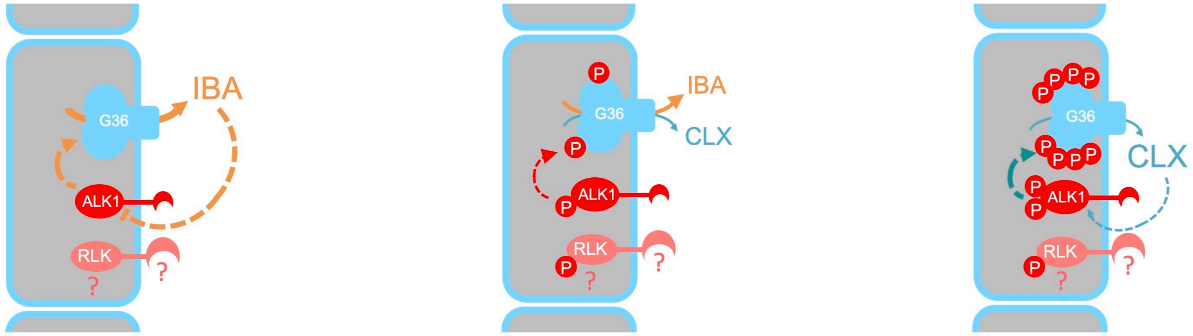


growth state:
dephosphorylation

default state:
intermediate phosporylation

defence state:
phosphorylation:

**Supplementary Figure 9: Hypothetical model on the action of ALK1/QSK1/KIN7 on ABCG36 phosphorylation**

In a default state (middle panel), ABCG36 (blue) is phosphorylated on an intermediate level by the LRR-RLK, ALK1/QSK1/KIN7 (dark red), allowing for IBA and CLX export, although IBA export might be preferred due to its higher affinity toward ABCG36. ALK1/QSK1/KIN7 acts like a classical co-receptor in an interaction with a not yet identified receptor-like kinase (RLK; light red)). In a growth state (left panel), ALK1/QSK1/KIN7 phosphorylation is reduced by exported IBA leading to activation of ABCB36-mediated IBA (but not CLX) export. Contrarily, in a defense state (right panel) CLX would promote ALK1/QSK1/KIN7 and thus ABCG36 phosphorylation leading to inhibition of IBA export, which allows for CLX export. Note that export of both ABCG36 substrates, IBA and CLX, would lead in this model to a self-amplification of growth and defense states, respectively. Straight lines indicate transport, while dashed lines refer to a signaling function including protein phosphorylation.

### Supplementary Tables

**Supplementary Table 1: ABCB36/PEN3/PDR8- interacting proteins identified by co-immunoprecipitation using ABCG36-GFP (PEN3:PEN3-GFP) lines followed by LC-MS/MS.**
Three independent co-immunoprecipitation/MS-MS analyses identified a short list of 62 common, putative ABCG36 interacting proteins that are sorted by the average number of significant sequences (n=3). ABCG-type transporters are coloured in blue, receptor-like kinases (RLKs) in red; ABCG36/PEN3/PDR (At1g59870) and ALK1/QSK1/KIN7 (At3g02880) are in bold.

**Supplementary Table 2: ALK1-interacting proteins identified by co-immunoprecipitation using ALK1-YFP as a bait followed by LC-MS/MS analyses.**
ABCG and receptor-like kinase (RLK) proteins relevant for this study are marked in red and blue, respectively; ABCG36 and ALK1 are marked additionally in bold. Prior to trypsin digest, 10 bands were size-selected by silver stain and manually cut out of the gel (fraction 1-10).

**Supplementary Table 3: Quality control of phospho-proteomics**
Abundance of indicated ABCG36 (blue) and ALK1 (red) phospho-sites identified by label-free phospho-proteomics of Arabidopsis Wt (Col Wt) or *alk1-2* grown in liquid cultures. Wt treated with IBA, CLX or FOX elicitor (1 µM, 24h) or infected with *F. oxysporum* (107 conidia/ml; 24h).

| target name | acc. number | primer sequences |
|---|---|---|
| *ABCG36* | *At1g59870* | fw: CAT GGA CCG TGT ATG GAG TG<br>rev: AGA CGG TGA AAG CGA TGA GTG |
| *ABCG37* | *At3g53480* | fw: TTG CGA TGT TCC TCG TCT C<br>rev: GAG TGT CCA AGA CGT TGG TG |
| *ALK1* | *At3g02880* | fw: AAC CGT ATT GAT GGC TAC CG<br>rev: ACC CAT CTC GGC AAA TCT AC |
| *AHA2* | *At4g30190* | fw: GGC ACT TGC TCA AAG GAC AC<br>rev: GCT TCA CGA CTG ATT CCA C |
| *NIP5;1* | *At4g10380* | fw: ATT GGC AGG TAT AGC CGT TG<br>rev: GTA GAC CGC TGC ACC AGA TAT G |
| *MYB51* | *At1g18570* | fw: CTA CAA GTG TTT CCG TTG ACT CTG AA<br>rev: ACG AAA TTA TCG CAG TAC ATT ACA GGA |
| *WRKY11* | *At4g31550* | fw: CCC ACG TGG TTA CTA CAA GTG C<br>rev: TGG ATC ATC TAA TGC TCG TTC CAC |

**Supplementary Table 4: Forward (fw) and reverse (rev) primers used for real-time PCR analyses in this study (see Suppl. Fig. 2B).**
